# Thermal biology of aphids and implications for agriculture and food security

**DOI:** 10.1101/2024.10.13.618050

**Authors:** Oswaldo C. Villena, Ali Arab, Peter A. Armbruster, Elizabeth Markovich, Kevin Zhang, Henri E.Z. Tonnang, Bonoukpoè M. Sokame, Jan Kreuze, Pablo Carhuapoma, Heidy Gamarra, Leah R. Johnson

## Abstract

Climate change poses significant challenges to agriculture and food security, particularly through its effects on insect vector populations and the pathogens they transmit. Aphids are one of the biggest group of ectotherms that transmit viruses to plants; more than 200 species have been identified as pathogen vectors. These aphids are responsible for transmitting over 300 viruses. The life-history traits of aphids such as fecundity and survival respond strongly and non-linearly to temperature and therefore to global warming. In this study, we elaborate the thermal responses for the main life-history traits (i.e, development and mortality rate) for aphid species for which data were available or generated. Also, thermal responses for virus transmission rates were elaborated to describe plant host to vector and vector to plant host dynamics. With these data, we elaborated thermal suitability models which were used to map current and projected scenarios for the transmission of viruses by aphids. Data was only available/generated for 19 aphid species, many of which only have data either at the lower or upper thermal limits. For virus transmission rates, from host plant to vector and from vector to plant data was only available/generated for potato virus Y (PVY), potato virus A (PVA), and potato leaf roll virus (PLRV) transmitted by the aphid *Myzus persicae*. Projections show that virus transmission by aphids will shift to northern latitudes. Understanding the thermal biology of aphids is crucial for developing effective measures to safeguard agriculture, especially staple crops, and food security in the context of climate change.

## Introduction

Arthropod vectors that transmit pathogens are detrimental to human well-being because they cause multiple diseases in humans, animals, and plants. Globally, vector-borne diseases are responsible for approximately 700,000 human deaths annually (Rocklöv and Dubrow, 2020) and infections in animals and crops of economic importance cause significant economic losses (Narladkar, 2018; Sundstrom et al., 2014). Pathogens transmission is tightly linked to the ecology and biology of vector populations which depends strongly on climate, especially temperature (Singh et al., 2023).

Climate change is already affecting crop yields through changes in temperature and precipitation patterns. The yields of top global crops (corn, wheat, rice, and potatoes), which provide most of human caloric intake, have decreased over the last 15 years (Adhikari et al., 2015; Gammans et al., 2017; Ray et al., 2019). Lower yields reduce global food supplies and increase the number of people experiencing food insecurity (Mbow et al., 2020). Climate change also influences the occurrence, prevalence, and severity of crop pests and diseases (Luck et al., 2011). When crop pests and disease outbreaks occur, the impact can be devastating. For example, the Irish potato famine caused by the fungal disease late blight (*Phytophthora infestans*) killed around one million people and a million more emigrated (Kynealy, 1995). More recently, invasions of desert locust (*Schistocerca gregaria*) throughout the Horn of Africa have shown how vulnerable crops are to pests (Salih et al., 2020).

As a result of climate change, temperature has been increasing in the last century. Global warming is projected to alter the distribution of crop pests/diseases, especially of ectotherms (Freeman et al., 2018). Traits of crop pests/diseases (e.g., development rate, survival rate, fecundity) are highly temperature sensitive (Sinclair et al., 2016). Extensive research has established that most species’ physiological and life-history traits respond non-linearly to changing temperatures, with performance increasing from zero at a thermal minimum to a high level at an optimum temperature before declining back to zero at a thermal maximum (Dell et al., 2011). Within the range of permissive temperatures, the non-linear influence of temperature on crop pest and disease traits affects the population size, the geographic distribution, and the magnitude of pathogens transmission (Bebber et al., 2014). Determining the effects of temperature on pathogens transmission requires identifying temperatures that determine the tradeoffs between different temperature-dependent crop pest and disease traits. In the last decade, there have been major improvements in mechanistic disease models for assessing the effect of temperature on vector-borne diseases such as malaria and dengue using life-history traits of vectors and parasites/viruses they transmit. These models allow investigators to estimate rates of transmission and to delineate current and future trends of vectors and parasites they transmit (Johnson et al., 2015; Mordecai et al., 2019; Ryan et al., 2023; Villena et al., 2022). Here, we develop a similar approach to assess the impact of changes of temperature patterns on pests (e.g., aphids, whiteflies) and pathogens (e.g., virus, bacteria) they transmit that cause diseases on staple crops and crops of economic importance.

Multiple important gaps limit our understanding about the effects of temperature on insect-borne diseases in staple crops and crops of economic importance (Singh et al., 2023). In this study, we summarize scientific knowledge about the effect of temperature on aphid-borne disease transmission, identify critical knowledge gaps, and discuss priorities for future research in the context of global warming and climate change. Aphids are the major insect group responsible for the transmission of viruses to crops. First, we aggregate available and generated data for life-history traits of aphids as well as virus transmission rates from vector to plants and vice versa. Second, we develop mechanistic thermal performance curves for each life-history trait and for virus transmission rates; we then use these performance curves to estimate a thermal suitability metric S(T) for the systems vector/pathogen for which data were available. Next, using the suitability models we elaborate thermal suitability maps (i.e., where the probability of S(T) > 0 is at least 0.95) for current and projected scenarios.

This study provides an important foundation for developing strategies to mitigate the effects of global warming on agriculture diseases by: 1) projecting changes in temporal and spatial components of aphid-transmitted crop disease risk in response to global warming, 2) developing a platform for developing similar models in other vector-pathogen-crop systems, and 3) compiling a comprehensive data set of the temperature sensitivity of aphid life-history and virus transmission rates that can be used as a basis for additional analysis and to identify current gaps in data availability.

## Materials and Methods

### Thermal trait data acquisition

An exhaustive literature review was performed to identify studies quantifying how temperature affects the life-history traits of aphid species that transmit pathogens to staple crops (e.g., potatoes, maize, wheat). We focused on the following traits: development time, development rate, fecundity (daily and total), reproduction rate, proportion nymphs that reach adulthood, longevity, oviposition duration, aphid size, and mortality rate. Also, we collected data on virus transmission rates from host plants to vectors, virus transmission rates from vectors to host plants, and viral titers. Definitions of each trait are in Table 1. To make direct comparisons amongst different temperatures, studies with fluctuating temperature were not included.

**Table 1.** Aphid life-history traits and definitions.

| **Trait** | **Trait Definition** | **Trait Name (Abbrev)** | **Trait Unit** |
| --- | --- | --- | --- |
| Development time | Mean development time at temperature $x$ | adt | days |
| Development rate | Rate at which the number of organisms in a population increases ($1/\text{development time}$) | adr | |
| Mortality nymphs | Proportion of death nymphs before adulthood at temperature $x$ | mun | percentage |
| Proportion nymphs to adulthood | Proportion of nymphs that reached adulthood | pna |  |
| Longevity | Mean days alive at temperature $x$ | lgv | days |
| Mortality rate | Rate of mortality for adults at temperature $x$ ($1/\text{longevity}$) | mu | |
| Fecundity (daily) | Mean number of nymphs per female per day at temperature $x$ | nfd | Individuals |
| Fecundity (total) | Mean number of nymphs per female per lifespan at temperature $x$ | nfl | Individuals |
| Transmission rate vector | % of individuals acquiring the virus | trv | percentage |
| Transmission rate host | % of individuals infected | trh | percentage |

Extracted data included the consumer (aphid), temperature, measured traits (with standard errors), study location (latitude and longitude), aphid stage, aphid resource (plant species), resource stage (e.g. leaf, twig), and pathogen (if applicable). The WebPlotDigitizer 4.6 software (Rohatgi, 2023) was used for data extraction if results were solely presented in the form of graphs or figures. In some cases, authors provided the raw data.

### Bayesian fitting of thermal traits

Most thermal traits (e.g., development rate, fecundity) of ectotherms (e.g., aphids, psyllids, mosquitoes) exhibit unimodal performance curves (Colinet et al., 2015; Mordecai et al., 2019). We fit each unimodal thermal response for all traits (Table 1) for each aphid species (e.g., *Myzus persicae, Aphis gossypii*) for all crops with available data (i.e., *Mizus persicae*/*Solanum tuberosum*). In cases where there were very few data points, we combined the data for the target aphid trait with different crops. In a similar way, we fit each unimodal thermal response for the transmission rate of the viruses transmitted by aphids (e.g., potato virus Y, potato virus A). To do this we used a Bayesian approach using the JAGS/*rjags v 4-15* package (Plummer et al., 2023) in R (R Development Core Team, 2017).

We defined an appropriate likelihood for each trait (e.g., binomial likelihoods for proportion data, truncated normal for continuous numeric traits) with the mean defined by a Briére function (Briere et al., 1999) for asymmetric responses like aphid development rate using the function:

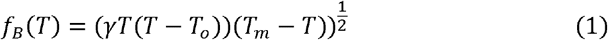

where *T*_0_ is the lower thermal limit (where the response becomes zero), *T*_*m*_ is the upper thermal limit, and γ is a constant that determines the curvature at the optimum. Formally we assume a piecewise continuous function, so that the thermal trait is assumed to be zero if *T* < *T*_0_ or *T* > *T*_*m*_.

For symmetric responses we used either a quadratic concave down or concave up function (Amarasekare and Savage, 2012; Villena et al., 2022). For concave-down symmetric responses like fecundity and the proportion of nymphs reaching adulthood, we fit a quadratic function parameterized in terms of the temperature intercepts, using the function:

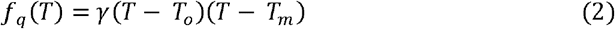

where *T*_0_ is the lower thermal limit, *T*_*m*_ is the upper thermal limit, and γ is a constant that determines the curvature at the optimum. As with the Briére function we assume the trait is piecewise zero above and below the thermal limits. For concave-up symmetric responses like mortality rate, we fit a concave-up quadratic function (Mordecai et al., 2019; Villena et al., 2022) using the function:

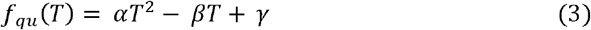

where α, β, and γ are the standard quadratic parameters. Note that because all traits must be ≥ 0 we also truncated this function, creating a piecewise continuous function where *f*_*qu*_ is set to zero if the quadratic evaluates to a negative value.

For all traits we chose relatively uninformative priors that limit each parameter to its biologically realistic range. More specifically, we assumed that temperatures below 0°C and above 45°C are lethal for aphid species or that their physiological functions cannot be performed (e.g., reproduction). Same lethal temperatures apply to the pathogens they transmit (Dampc et al., 2021; Hazell et al., 2010). Based on these assumptions we set uniform priors for the minimum temperature (*T*_0_) between 0 – 24°C and for the maximum temperature (*T*_*m*_) between 25 – 45°C (Dampc et al., 2021; Taylor et al., 2019). Priors for other parameters in the thermal responses were set to ensure parameters were positive and not tightly constrained.

The *rjags* package uses a Metropolis algorithm within Gibbs Markov Chain Monte Carlo (MCMC) sampling scheme to obtain samples from the joint posterior distribution of parameters. For each fitted trait, we obtained posterior samples from five Markov chains that were run for 20000 iterations initiated with random starting values. These samples were obtained after using 10000 iterations for adaptation and burning another 10000 iterations. We visually assessed convergence of the Markov chains. To obtain the posterior summaries of each trait, we combined the 20000 samples from each Markov chain from the posterior distribution, which resulted in a total of 100000 posterior samples of the thermal response for each trait. Based on these samples, for each unimodal thermal response we calculated the posterior mean, the 95% highest probability density (HPD) interval, and the 95% prediction interval (PI) around the mean of the thermal response and various summaries (e.g., the thermal minimum or maximum).

### Bayesian fitting of the suitability metric for aphid/pathogen

Once the posterior samples for all thermal traits across temperature for each species were obtained, we used these samples to estimate the number of aphids in the population using the following equation:

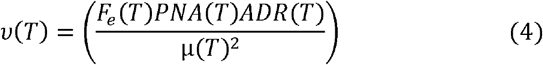

where FE is the total number of nymphs that a female will produce during its life cycle, PNA is the number of aphid nymphs that will reach adulthood, ADR is the aphid development rate, and is the aphid mortality rate (Taylor et al., 2019).

In a similar way we estimated the transmission rate from the insect vector, the aphid, to the host crop and from the host crop to the insect vector using the following equation:

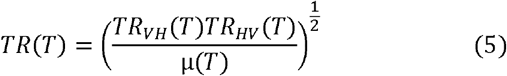

where *TR*_*V H*_ is the virus transmission rate from the aphid to the host plant, *TR*_*HV*_ is the virus transmission rate from the host plant to the aphid, and *µ* is the aphid mortality rate (Taylor et al., 2019).

Combining the number of aphids in the population (equation 4) and the transmission rates from vector to host plant and from host plant to vector (equation 5), we estimated a suitability metric *S(T)* across a temperature range which includes the posterior median and the 95% HPD of the overall thermal response of suitability, as well as the critical thermal minimum, thermal maximum, and optimal temperature for transmission suitability for the most completed data sets of our study, transmission rate of potato virus Y, potato virus A, and potato leafroll virus by *Myzus persicae* and *Aphis gossypii* using the following *S(T)* equation:

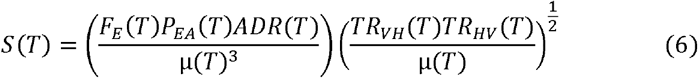

We also performed an uncertainty analysis for *S(T)* to quantify the contribution of each trait to the overall uncertainty in mean suitability.

### Mapping of the current and projected transmission thermal suitability in crops across space

For current transmission thermal suitability, we mapped the number of months a year that locations are predicted to have suitable temperatures for the transmission (risk of transmission) of viruses (e.g., potato virus A, potato leaf roll virus) by aphids (e.g. *Aphis gossypii*) for virus/vector subsystems for which data is available. For temperature data, we used monthly mean temperature rasters at a 5-minute spatial resolution downloaded from the WorldClim-Global Climate project (Fick and Hijmans, 2017). Risk maps were produced using the *mappestRisk* package in *R* (San-Segundo Molina et al., 2024). The maps only show temperatures that are highly suitable for the transmission of viruses by aphids, but they do not indicate transmission magnitude or intensity.

For projected transmission thermal suitability, we mapped future scenarios for the transmission of the potato leaf roll virus (PLRV) and the potato virus A (PVA) by the aphid *Aphis gossypii*. We assessed changes in year-round transmission for years 2040 and 2100 using two General Circulation Models (GCMs), the Mediterranean Centre on Climate Change Earth System Model 2 (CMCC-ESM2) from the Euro-Mediterranean Centre on Climate Change and the Hadley General Circulation Model (HadGEM3-GC31-LL) from the Hadley Centre for Climate Science and Services under two Shared Socioeconomic Pathways (SPP) scenarios: SSP126 and SSP585 (Ryan et al., 2023). The SSP126 scenario represents an optimistic scenario simulating a development that is compatible with the 2°C target and assumes climate protection measures being taken while the SPP585 scenario represents the upper boundary of the range of scenarios and assumes a fossil-fueled development. Risk maps and differences between scenarios were produced using the *mappestRisk* package in R (San-Segundo Molina et al., 2024).

## Results

### Thermal response of life-history traits and virus transmission rates

In this study, we aggregated data for life-history traits (Table 1) in response to temperature for 19 aphid species: *Acyrthosiphon kondoi* (blue alfalfa aphid), *Acyrthosiphon pisum* (pea aphid), *Aphis citricola* (brown citrus aphid), *Aphis fabae* (black bean aphid), *Aphis gossypii* (cotton aphid), *Aphis nasturtii* (damson-hop aphid), *Aulacorthum solani* (foxglove aphid), *Brachycaudus schwartzi* (peach curl aphid), *Brevicoryne brassicae* (cabbage aphid), *Erisoma lanigerum* (woolly apple aphid), *Hyperomyzus lactucae* (lettuce aphid), *Macrosiphum euphorbiae* (potato aphid), *Melanaphis sacchari* (sugarcane aphid), *Myzus persicae* (green peach aphid), *Rhopalosiphum maidis* (corn leaf aphid), *Rhopalosiphum padi* (bird cherry-oat aphid), *Schizaphis graminum* (greenbug), *Sitobion avenae* (English grain aphid), and *Toxoptera citricida* (brown citrus aphid) (Figures 1-3 and S1-S3). For virus transmission rates our study only identified data for the transmission of potato virus Y strain O (PVY-O) and potato virus A (PVA) from *Nicotiana benthamiana* (benthi) to *Myzus persicae* and potato leaf roll virus (PLRV) from *Physalis floridana* (garden berries) to *M. persicae* (Figure S4). For the virus transmission rates from *Myzus persicae* (the vector) to host plant, we only identified data for the transmission of PVY with *N. benthamiana, N. tabacum* (tabacco), and *Solanum tuberosum* (potatoes); PVY-O with *N. benthamiana* and *S. tuberosum*; PVA with *N. benthamiana*; and PLRV with *S. tuberosum* and *P. floridana* (Figure S5).

**Figure 1.**
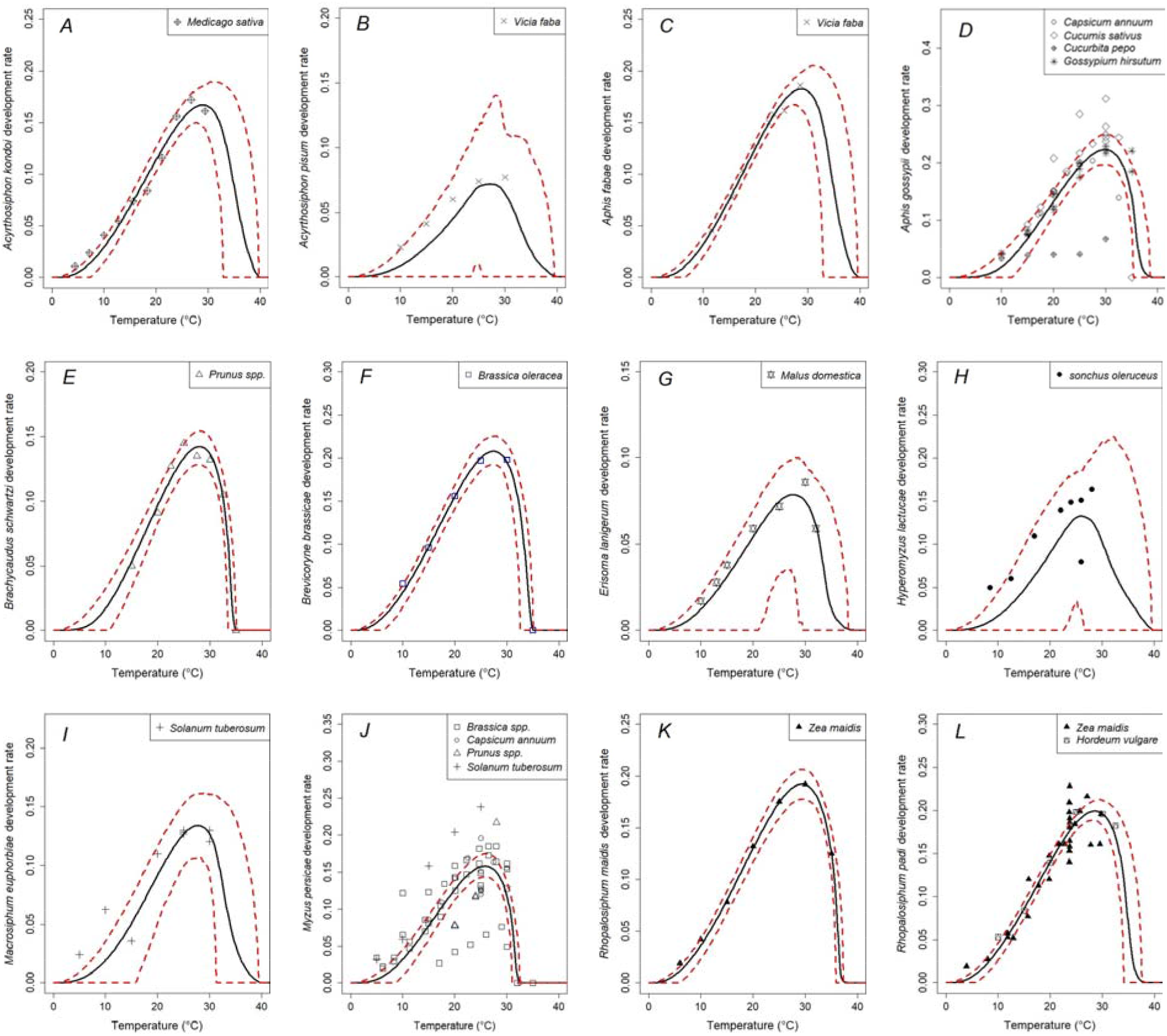
Posterior mean (solid line) and 95% highest posterior density (HPD; dashed lines) of the thermal responses for aphids development rate (ADR) for: A) *Acyrthosiphon kondoi*, B) *Acyrtosiphon pisum*, C) *Aphis fabae*, D)*Aphis gossypii*, D) *Brachycaudus schwartzi*, E) *Brevicoryne brassicae*, F) *Erisoma lanigerum*, G) *Hyperomyzus lactucae*, H) *Macrosiphum euphorbiae*, I) *Myzus persicae*, J) *Rophalosiphum maidis*, and K) *Rophalosiphum padi*.

### Posterior distribution of thermal traits

We estimated life-history trait posterior means and the 95% highest posterior density (HPD) of the thermal responses for 19 aphid species for which data was available or generated. For aphid development rate (ADR), we estimated the posterior mean and the 95% of the HPD for 12 species (Figure 1, Table 2). For aphid fecundity, the number of nymphs per female per lifecycle (NFL), we estimated the posterior mean and HPD for 16 species (Figure S1, Table S1). For the proportion of aphid nymphs surviving to adulthood (PNA), we estimated the posterior mean and HPD for 12 species (Figure S2, Table S2). For aphid mortality rate (*µ*), we estimated the posterior mean and HPD for 16 species (Figure S3, Table S3). For virus transmission rates from host plants to the aphid vector, data was only available for the transmission of potato virus Y, strain O (PVY-O), potato virus A (PVA), and potato leaf roll virus (PLRV) by *Myzus persicae* (Figure S4). For virus transmission rates from the aphid vector to the host plant, data was only available for the transmission of potato virus Y (PVY), potato virus Y, strain O (PVY-O), potato virus A (PVA), and potato leaf roll virus (PLRV) by *Myzus persicae* (Figure S5).

**Table 2.** Average thermal optima, thermal limits, and 95% confidence intervals (Cis) for aphid development rate.

| Species | Crops | Lower thermal<br>(CIs) | Optimum<br>Temp (CIs) | Upper thermal<br>(CIs) |
| --- | --- | --- | --- | --- |
| *Acyrtosiphon kondoi* | *Medicago sativa* | 12.6<br>(11.8, 13.4) | 29.2<br>(28.59, 29.81) | 35.05<br>(34.77, 35.33) |
| *Acyrtosiphon pisum* | *Vicia faba* | 14.55<br>(13.85, 15.25) | 27.9<br>(27.35, 28.45) | 32.55<br>(32.29, 32.81) |
| *Aphis fabae* | *Vicia faba* | 12.55<br>(11.76, 13.34) | 29.2<br>(28.59, 29.81) | 34.85<br>(34.57, 35.13) |
| *Aphis gossypii* | *Capsicum annuum*,<br>*Cucumis sativus*,<br>*Cucurbita pepo*,<br>*Gossypium hirsutum* | 15.25<br>(14.52, 15.98) | 30.0<br>(29.41, 30.59) | 35.3<br>(35.2, 35.4) |
| *Brachycaudus schwartzi* | *Prunus persica* | 14.25<br>(13.54, 14.96) | 27.9<br>(27.33, 28.47) | 33.2<br>(32.93, 33.47) |
| *Brevicoryne brassicae* | *Brassica oleracea* | 11.45<br>(10.66, 12.24) | 27.6<br>(27.01, 28.19) | 33.1<br>(32.82, 33.38) |
| *Erisoma lanigerum* | *Malus domestica* | 13.1<br>(12.35, 13.85) | 28.3<br>(27.72, 28.88) | 33.25<br>(32.98, 33.52) |
| *Hyperomyzus lactucae* | *Sonchus oleruceus* | 14.4<br>(13.73, 15.07) | 26.9<br>(26.37, 27.43) | 31.35<br>(31.08, 31.62) |
| *Macrosiphum euphorbiae* | *Solanum tuberosum* | 13.3<br>(12.55, 14.05) | 27.8<br>(27.23, 28.37) | 33.25<br>(32.98, 33.52) |
| *Myzus persicae* | *Brassica campestris*,<br>*Brassica oleracea*,<br>*Brassica rapa*,<br>*Capsicum annuum*,<br>*Prunus armeniaca*,<br>*Prunus persica*,<br>*Solanum tuberosum* | 12.1<br>(11.33, 12.77) | 25.8<br>(25.24, 26.36) | 30.85<br>(30.59, 31.11) |
| *Rhopalosiphum maidis* | *Zea mays* | 12.1<br>(11.28, 12.92) | 29.5<br>(28.89, 30.11) | 35.4<br>(35.07, 35.63) |
| *Rhopalosiphum padi* | *Hordeum vulgare*,<br>*Zea mays* | 12.3<br>(11.51, 13.09) | 28.8<br>(28.2, 29.4) | 34.3<br>(34.03, 34.57) |

We observed variability of thermal sensitivity between and within life-history traits. Aphid development rate has a higher average optimum temperature (28.24°C; CI: 25.8 - 30°C) compared to total fecundity (19.24°C, CI: 16.20 - 21.4°C), the proportion of nymphs reaching adulthood (20.06°C; CI: 18.4 - 23.2°C), and mortality rate (17.83°C; CI: 12.9 - 22.5°C) (Figure 2). Within life-history traits we also observed significant differences between aphid species. For aphid development rate, *Rhopalosiphum maidis* (12.1 - 35.4°C), *Acyrthosiphon kondoi* (12.6 - 35.05°C), and *Aphis fabae* (12.5 - 34.8°C) shows the largest tolerance thermal range while *Hyperomyzus lactucae* (14.4 - 31.35°C), *Acyrthosiphon pisum* (14.5 - 32.5°C), and *Myzus persicae* (12.1 - 30.8°C) show the shortest tolerance thermal range for ADR (Figure 1, Table 2). For fecundity, Aphids that produced the greater number of nymphs per female per lifecycle are *Erisoma lanigerum* (108 nymphs/female), *Acyrtosiphum kondoi* (87 nymphs/female), *Aulacorthum solani* (72 nymphs/female), and *Acyrtosiphum pisum* (70 nymphs/female). Furthermore, *Myzus persicae* and *Rophalosiphum maidis* are the aphids that showed the largest tolerance thermal ranges for nymph production (9.5 - 33.5°C and 10.85 - 29.95°C respectively) while *Aulacrothum solani* and *brachycaudus schwartzi* showed the shortest tolerance thermal ranges (13.25 - 24.3°C and 10.4 - 25.6°C respectively) (Figure S1; Table S1).

**Figure 2.**
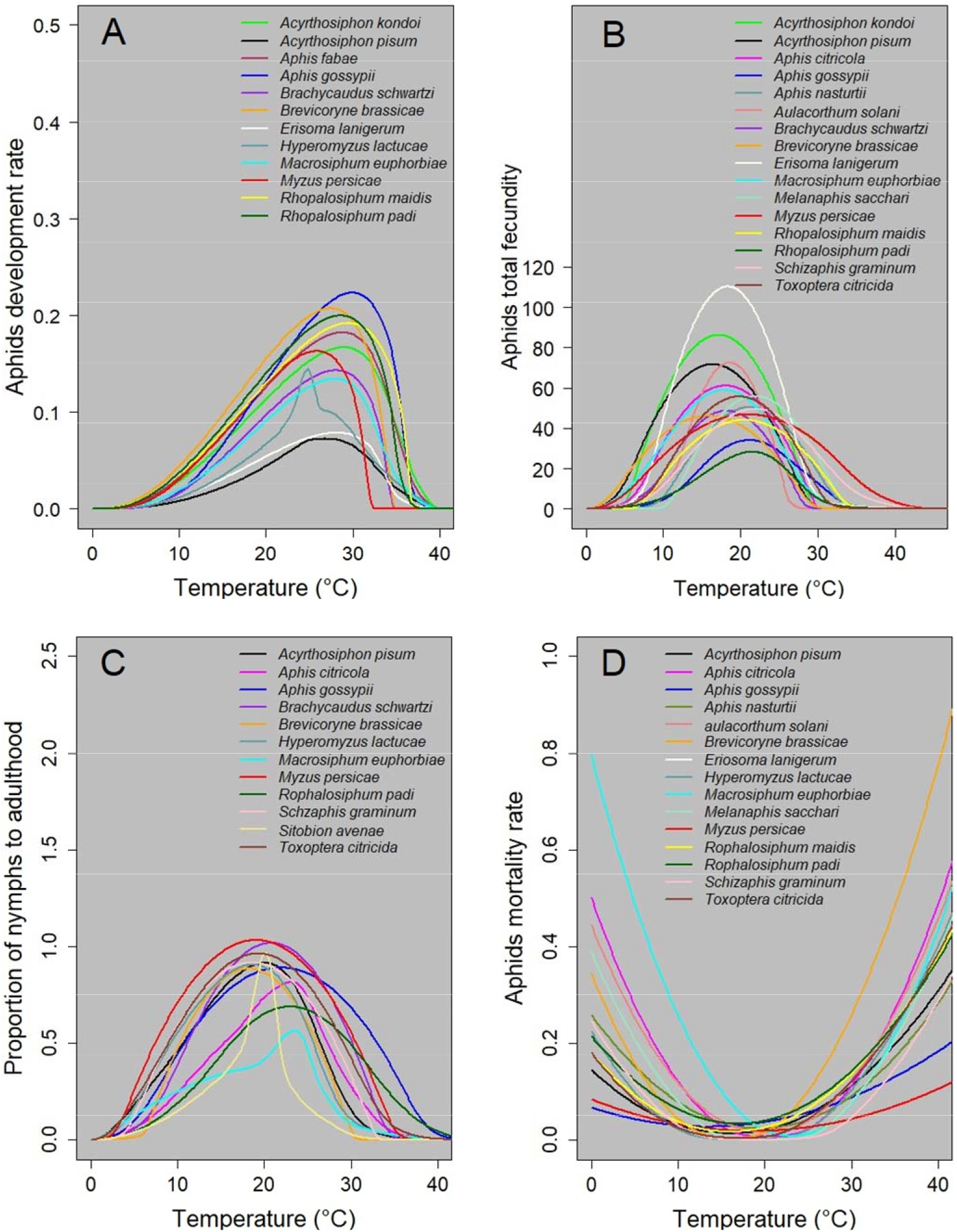
Aphids’ life-history traits differing thermal optima and thermal limits for: A) Aphid development rate (ADR), B) Aphid total fecundity (NFL), C) Aphid proportion of nymphs to adulthood (PNA), and D) aphid mortality rate (*µ*).

Similarly, for the proportion of nymphs reaching adulthood (PNA), *Aphis gossypii, Myzus persicae*, and *Aphis citricola* have the greatest tolerance thermal ranges with values of 8.6 - 35.15°C, 7.3 - 30.8°C, and 9.05 - 30.8°C respectively (Figure S2). On the other hand, the aphids with the lowest tolerance thermal range for PNA are *S. avenae, A. pisum*, and *M. euphorbie* with ranges of 15.4 - 23.45°C, 13.8 - 26.8°C, and 11.8 - 25.4°C respectively (Figure S2, Table S2). For mortality rate (*mu*), we observed different average optimum temperature for the lowest mortality rates. For example, *M. euphobuae* and *A. solani* showed lowest mortality rates when temperatures are 22.5 and 20.4°C respectively, while *A. gossypii* and *A. pisum* showed lowest mortality rates when temperatures are 12.9 and 15.7°C respectively (Figure S3; Table S3).

We also observed that thermal performance curves of aphid species are very similar between some crops while with other crops are slightly different. For example, *Aphids gossypii* development rate is very similar between *Capsicum annuum* and *Gossypium hirsutum*. However, *A. gossypii* development rate seems to be higher in *Cucumis sativum* but lower with *Cucurbita pepo* (Figure 1D). *Rhopalosiphum padi* development rate seems to be very similar to *Zea mays* and *Hordeum vulgare* (Figure 1L). Other examples are fecundity of *Toxoptera citricola* with *Citrus unshiu* and *Citrus aurantium* (Figure S1P), the upper thermal limit of *Macrosiphum euphorbiae* with *Lactuca sativa* and *Solanum tuberosum* (Figure S1J). Also, the proportion of nymphs to adulthood for *Aphis gossypii* with *Capsicum annuum, Citrus unshiu, Cucumis sativus*, and *Gossypium hirsutum* (Figure S2C); for *Myzus persicae* with *brassica spp, Capsicum annuum*, and *Prunus spp*. (Figure S2H); for *T. citricola* with *C. unshiu* and *C. aurantium* (Figure S2L).

### Posterior distributions of the thermal responses of virus transmissions

For the virus transmission rates from the host plant to the insect vector, there is only thermal response information for the transmission of potato virus Y, strain O (PVY-O), potato virus A (PVA) from *Nicotiana benthamiana* to *Myzus persicae* and potato leaf roll virus (PLRV) from *Physalis floridana* to *M. persicae* (Figure 3 and S4). At optimum temperature, PLRV virus transmission from plant host (*P. floridana*) to *M. persicae* is higher (83%) compared to the transmission of PVA virus (64%) and PVY-O (38%) from *N. benthamiana* to *M. persicae* (Figure S4).

**Figure 3.**
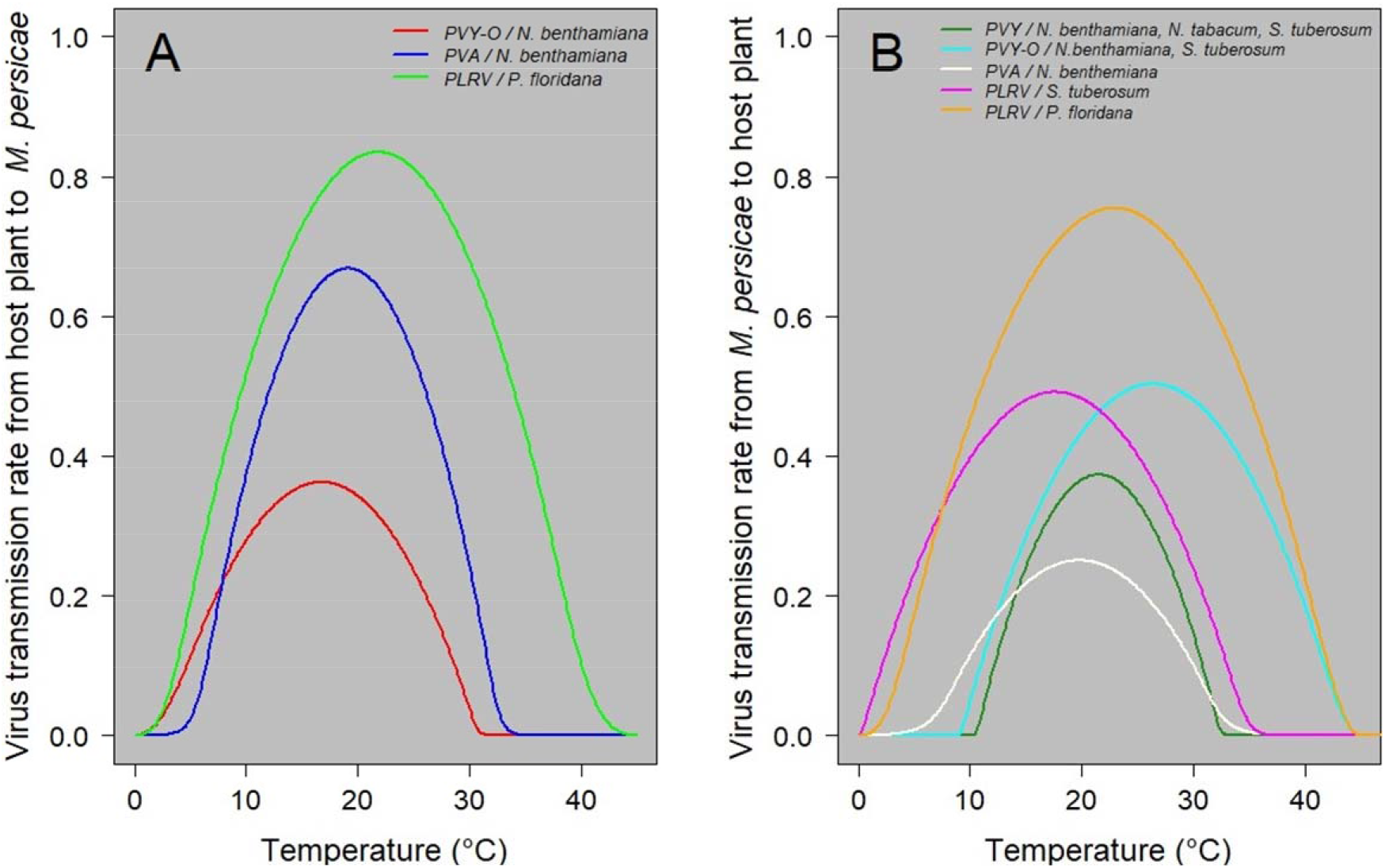
Thermal response of virus transmission rates from: A) host plant to *Myzus persicae*, B) *Myzus persicae* to host plant. Viruses: potato virus Y, strain O (PVY-O), potato virus A (PVA), potato leaf roll virus (PLRV). Crops: *Nicotiana benthemiana, Physalis floridana, Solanum tuberosum*, and *Nicotiana tabacum*.

For the virus transmission rates from the insect vector to the host plant, there is only information for PLRV from *M. persicae* to *solanum tuberosum*, PLRV from *M. persicae* to *P. floridana*, PVY from *M. persicae* to *N. benthamiana, N. tabacum*, and *Solanum tuberosum*, PVY-O from *M. persicae* to *N. benthamiana* and *S. tuberosum*, and PVA from *M. persicae* to *N. benthamiana* (Figure 3 and S5). There is higher transmission rate for PLRV (78%) from *M. persicae* to host plant (*P. floridana*). Next higher transmission rates are for PVY-O (51%) and PLRV (50.5%) from *M. persicae* to *N. benthamiana* and *S. tuberosum* (Figure S5).

### Posterior distribution of S(T)

We estimated the posterior median and 95% HPD of the S(T) for the transmission of potato virus A (PVA) and potato leaf roll virus (PLRV) by *Aphis gossypii* because data for these virus/aphid systems are the most complete. Transmission of PVA and PLRV by *A. gossypii* have similar lower thermal limits (13.62°C and 13.65°C respectively) and optimum temperature of transmission (19.28°C and 19.39°C respectively). The main difference is in the upper thermal limit. For the transmission of PVA, the upper thermal limit is 32.59°C while for PLRV it is 34.61°C (Figure 4). However, these median ranges mask a great deal of uncertainty which could be reduced as more data become available.

**Figure 4.**
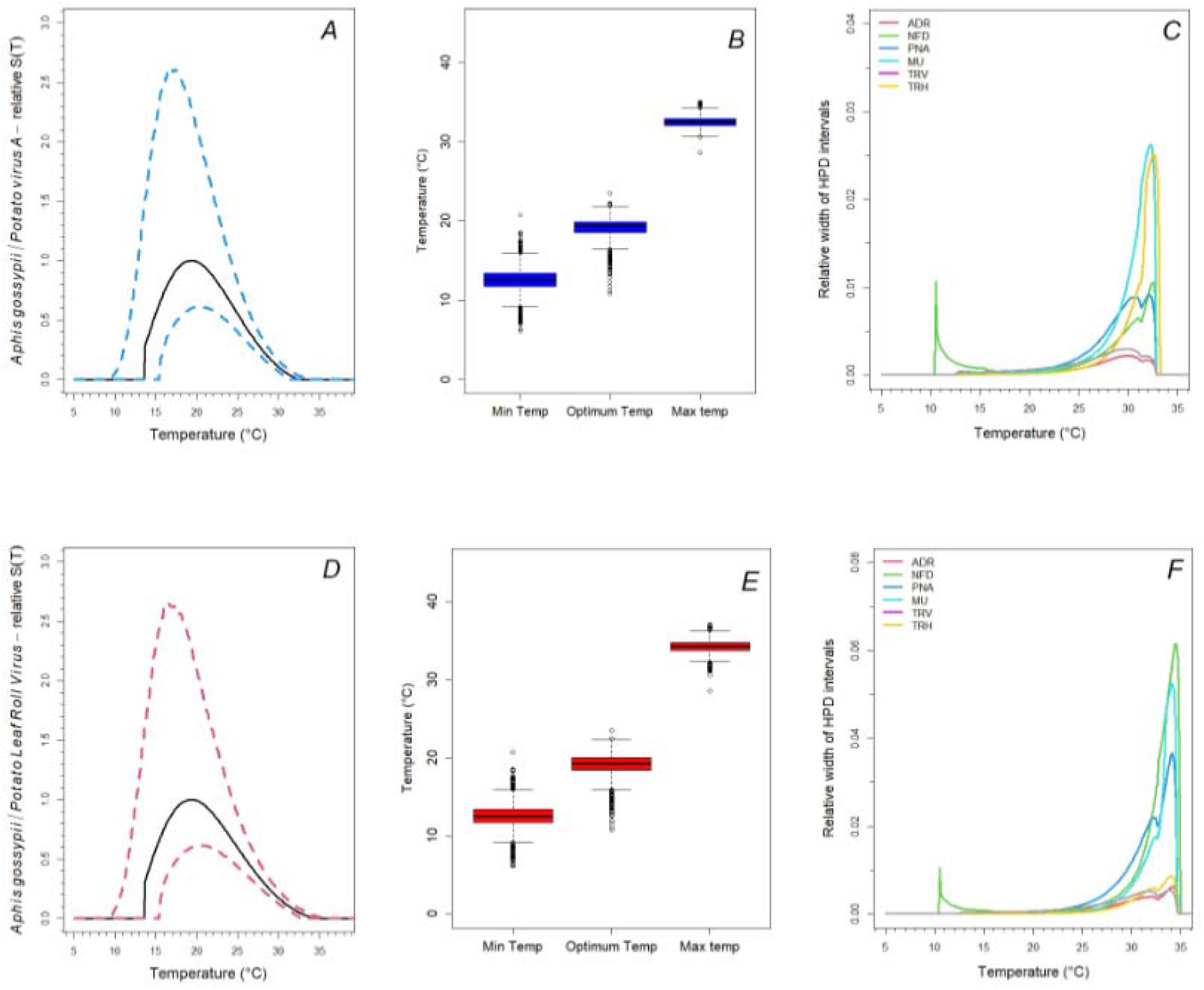
Top row: Relative transmission suitability S(T) for potato virus A (PVA) transmitted by *Aphis gossypii*. Bottom row: Relative transmission suitability s(T) for potato leaf roll virus (PLRV) transmitted by *Aphis gossypii*. A,D) Relative transmission suitability S(T) and 95% HPD, B,E) median, interquartile range, minimum and maximum numbers from the posterior for the minimum, optimum, and maximum temperatures, C,F) Relative width of the 95% HPD intervals due to uncertainty in each component, compared to uncertainty in S(T) overall.

Uncertainty for the transmission of PVA by *A. gossypii* is driven by daily fecundity at the lower thermal limit while mortality rate and the virus transmission rate from vector to host drives uncertainty in the upper thermal limit (Figure 4C). Uncertainty for the transmission of PLRV by *A. gossypii* is driven by daily fecundity at the lower thermal limit while daily fecundity, mortality rate, and the proportion of nymphs reaching adulthood drives uncertainty in the upper thermal limit (Figure 4F).

### Mapping of current and projected climate suitability for the transmission of viruses to potato crops

Our maps show the number of months per year that locations fall within suitable temperatures at current times for the transmission of potato virus A (PVA) and potato leaf roll virus (PLRV) by *Aphis gossypii* (Figure 5) and the regions where potatoes are produced (Figure S8). In North America, some potato production regions on the west and east coast have 4 to 8 months of suitable temperatures for the transmission of PVA and PLRV by *A. gossypii* (Figure 5). In Central America, potato production regions in Mexico, Guatemala, Honduras, Nicaragua, and Panama have between 8 to 10 months of suitable temperatures for the transmission of PVA and PLRV by *A. gossypii* (Figure 5). In South America, potato production regions in Venezuela, Colombia, Ecuador, Peru, Bolivia, and Brazil have year-round suitable temperatures for the transmission of PVY and PLRV (Figure 5). In Africa, there are suitable temperatures from 10 months to year-round transmission of PVA and PVY in the potato production regions of Angola, Zimbabwe, Mozambique, Tanzania, Kenya, Ethiopia, Sudan, and east of Madagascar (Figure5).

**Figure 5.**
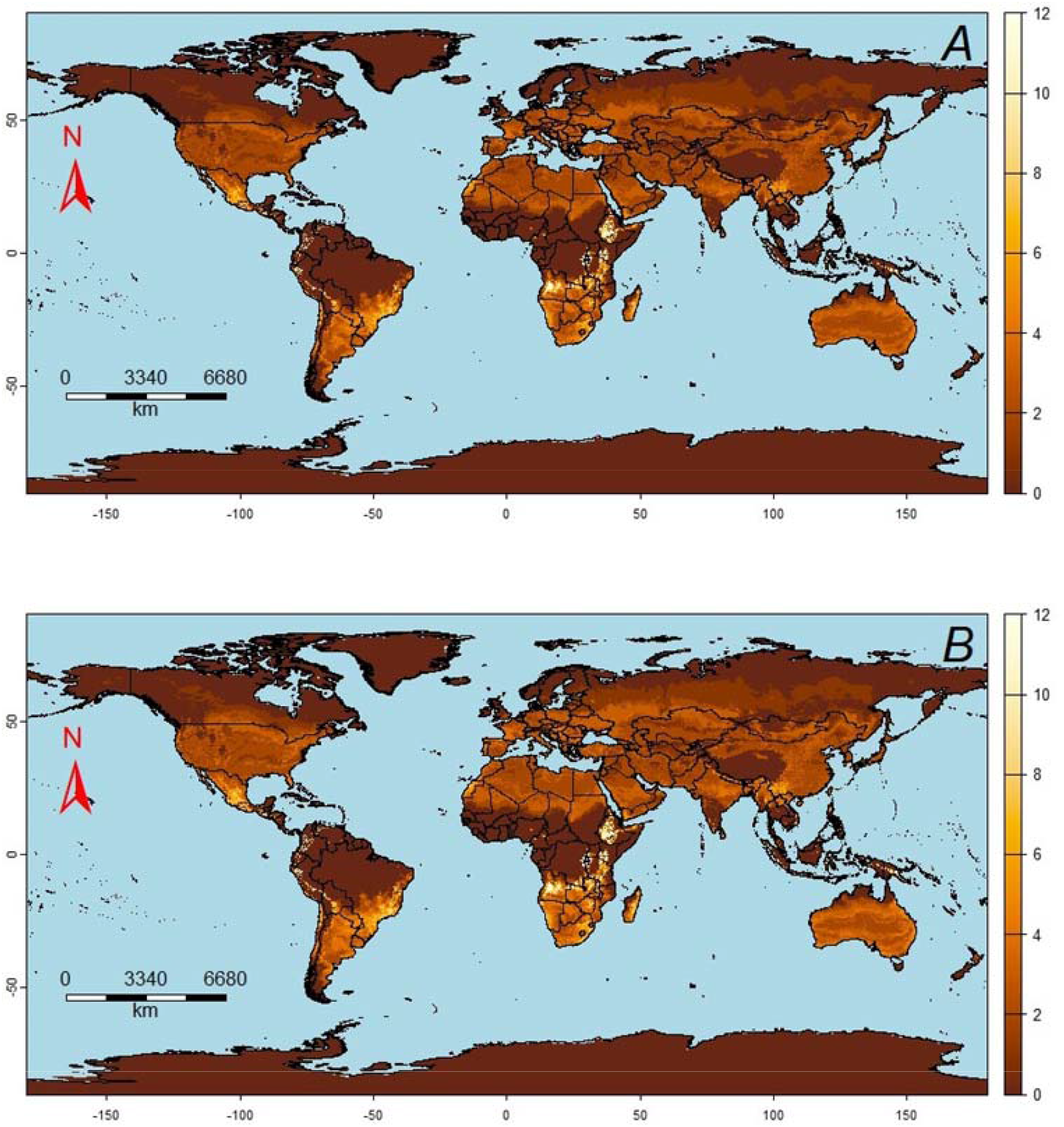
The number of months a year that locations fall within the predicted suitable range (probability of S(T) > 0 ≥ 0.95) for the transmission of: A) potato virus A (PVA) by *Aphis gossypii* and B) potato leaf roll virus (PLRV) by *Aphis gossypii*.

Predicted year-round transmission suitability for 2040 and 2100 for both models, HadGEM3-GC31-LL and CMCC-ESM2, for SPP126 and SPP585 scenarios are shown in Figures S9 and S11. The gains (increase in area) and losses (decrease of area) in year-round transmission between projected years and scenarios are shown in Figures 6 and S10. Both models show similar results. An expansion of year-round transmission suitability for PLRV by *A. gossypii* is seen in both scenarios SPP126 and SPP585. The potential for year-round transmission above the equator is predicted to expand at northern latitudes, where areas with no current year-round transmission will have the potential for transmission year-round. Below the equator, PLRV year-round transmission is predicted to expand in the potato production regions of Colombia, Ecuador, Peru, Bolivia, and south of Chile (Figures 6 and S10). There are no major differences between the SPP126 and SPP585 scenarios for 2040. There are only small increases in year-round transmission of PLRV in the potato production regions of Tamaulipas in Mexico and in the south of Chile and Argentina and in the coastal area of the states of Santa Catarina, Parana, and Sao Paulo in Brazil (Figures 6A and S10A). In the northern African region, Yemen, Oman, and Saudi Arabia there are more decreases than increases in areas with year-round transmissions of PLRV (Figures 6A and 10A).

**Figure 6.**
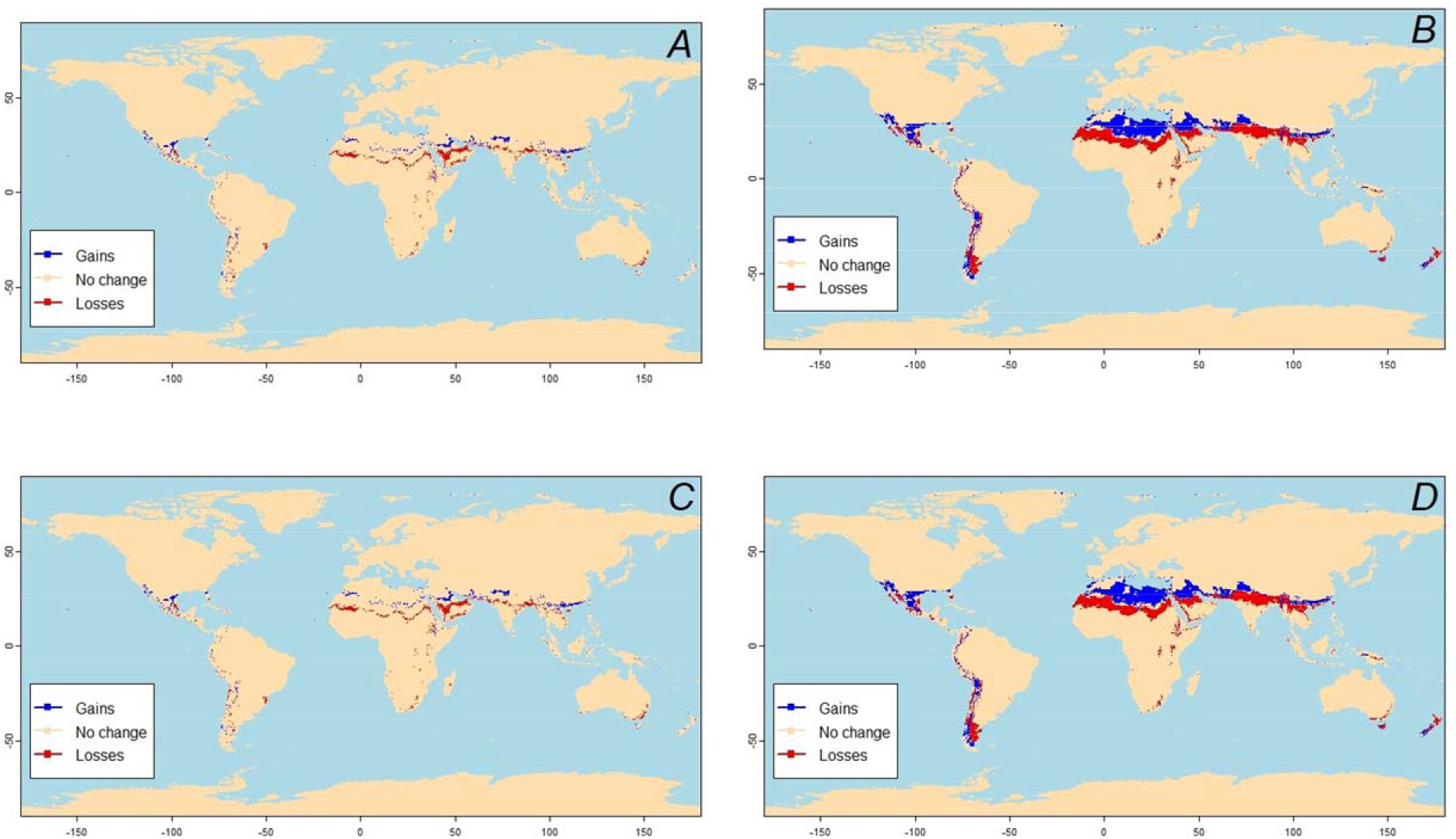
Changes in year-round thermal suitability projections for the transmission of potato leaf roll virus (PLRV) by *Aphis gossypii* using the HadGEM3-GC31-LL model for: A) changes at year 2040 between SSP126 and SSP585 scenarios, B) changes at year 2100 between SSP126 and SSP585 scenarios, C) changes between years 2040 and 2100 for SSP126 scenario, and D) changes between years 2040 and 2100 for SSP585 scenario. Gains mean new areas where PLRV virus can be transmitted year-round by *Aphis gossypii*; Losses mean areas where PLRV virus cannot be transmitted year-round anymore by *Aphis gossypii*, where the posterior probability of S(T) > 0 is 0.95.

For 2100, under the conservative scenario SSP126, in the Americas there will be an increase in areas with PLRV year-round transmission in most of Mexico potato production regions, in the northern of Florida, and in South America in Colombia, Ecuador, Peru, Bolivia, and south of Chile. In Africa, PLRV year-round transmission will increase in Morocco, Algeria, Libya, Egypt and very small increases in the potato production regions of Rwanda, Burundi, Kenya, and Malawi. In south-east Asia, there are some increases in the potato production regions of Saudi Arabia, Iran, Pakistan, Tibet, and the regions of Guangxi, Shenzhen, Guangdong, and Fujian in China. More of the decreases are in Yemen, Oman, and in India (Figures 6C and 10C). For the least conservative scenario SSP585, most of Mexico and the northern area of Florida will be suitable for PLRV year-round transmission. In South America, most of the potato production regions of Colombia, Ecuador, Peru, Bolivia, and southern region of Chile will be suitable for PLRV year-round transmission. In Africa, the potato production regions of Morocco, Tunisia, Algeria, Libya, and Egypt will have suitable temperatures for PLRV year-round transmission. In south-east Asia, most of the potato production regions of Saudi Arabia, Iran, Pakistan, Nepal, Tibet, and the regions of Nauning, Guangxi, Shenzhen, Guangdong, and Fujian in China will have suitable temperatures for PLRV year-round transmission. Also, most of the territory of Tasmania and New Zealand will be suitable for PLRV year-round transmission by *A. gossypii* (Figures 6D and S10D).

## Discussion

The continuous increase in global temperatures will undoubtedly affect the transmission of plant pathogens by insect vectors, which in turn will pose new threats to agriculture production and food security. Higher temperatures can affect insects’ growth rate, fecundity, phenology, and overall fitness. While some species can tolerate the impacts of higher temperatures, many insect and pathogen species will respond to higher temperatures by shifting their ranges poleward or to higher altitudes, by evolving adaptations to the changed temperatures, or by going extinct (Berg et al., 2010; Harvey et al., 2023). As temperatures rise, the biological and ecological dynamics of aphid populations will undergo significant changes as will aphid-mediated virus transmission, leading to increased risks for agriculture ecosystems globally and indirectly having broader implications for global food security (Ristaino et al., 2021; Singh et al., 2023).

The impact of global warming on virus transmission by aphids to staple crops and crops of economic importance is complex and profound. For example, modelling projections show that by 2050, between 2.5 to 4.2 billion people will be at risk of undernourishment because of population increase, climate change, lack of farmland, and crop yield reduction (Dawson et al., 2016; Watts et al., 2018). During this century, it is expected that earth’s average surface temperature will exceed the “safe” threshold of 2°C above pre-industrial average temperature. 2023 was already the warmest year since global record-keeping began in 1850 (Esper et al., 2024).

In recent years, multiple studies have concluded that climate change, conflict, and the Covid-19 pandemic are the main drivers of current high levels of global food insecurity. However, crop damage from agricultural pests and/or plant diseases are generally not taken into account in these studies, even though many of these plant diseases are greatly impacted by changes in climate. The lack of studies related to the impacts of climate change, especially global warming, on pest biology and/or pest vectored diseases in crops is surprising, considering the major pest outbreaks in recent years, outbreaks such as the fall army worm (Diaz-Alvarez et al., 2023; Rukundo et al., 2020; Tippannavar et al., 2019), maize lethal necrosis (Mahuku et al., 2015; Regassa et al., 2024), and the desert locust plagues that devastated crops in East Africa and parts of the Middle East and Asia (Salih et al., 2020).

Our study revealed a scarcity of data on how temperature impacts aphid species’ life-history traits (i.e. development rate, fecundity, mortality). For example, there is data for only 12, 16, 12, and 15 aphid species for development rate, fecundity, proportion of nymphs reaching adulthood, and mortality rate, respectively. Furthermore, uncertainty across the thermal performance curves for each life-history trait and for each aphid species is driven by data scarcity. For example, for *Acyrthosiphon pisum* and *Erisoma lanigerum* we only have 5 and 7 data points respectively and most of these points are in the lower thermal range (< 25°C). This is very similar to the situation with other aphid life-history traits. Only 6 of the 16 aphid species with available data for fecundity have more than 10 data points across our thermal range of study (0 to 40°C). Similarly, only 8 of 12 and 10 of 15 aphid species with available data for the proportion of nymphs that reach adulthood and mortality rate have more than 10 data points across the thermal range of our study. Furthermore, the thermal response of aphid’s life-history traits are only available for very few crops of economic importance.

The thermal range variation of aphid life-history traits (i.e., development rate, fecundity) among different aphid species highlights the need for more research to understand aphid population dynamics and their potential impact on agriculture and food security. For development rates, aphids exhibit an average lower thermal limit of 13.2°C, an optimum temperature of 28.2°C, and an upper thermal limit of 33.5°C with *R. maidis, A. kondoi*, and *A. fabae* showing broad thermal ranges, while *H. lactucae, A. pisum*, and *M. persicae* showing narrower thermal ranges. Similarly, fecundity exhibits an average lower thermal limit of 10.7°C, an optimum temperature of 19.24°C, and an upper thermal limit of 27.8°C with *E. lanigerum, A. kondoi*, and *A. pisum* showing high fecundity rates, *M. persicae* showing the widest thermal range (9.5°C to 33.5°C) for nymph production. The study of (Sinclair et al., 2012) highlights the temperature sensitivity of ectotherms, including aphids, emphasizing that their physiological and life-history traits respond non-linearly to temperature changes, affecting their development and reproduction rates. Understanding the underlying mechanisms of how temperature affects aphid life-history traits dynamics is important for pest management (Colinet et al., 2015).

The scarcity of data on the rates of virus transmission from aphid vectors to plant hosts and vice-versa represents an even more serious knowledge gap. Our study identified data for the transmission of potato virus Y, strain O (PVY-O), potato virus A (PVA), and potato leaf roll virus (PLRV) from host plants to only one aphid species, *Myzus persicae*. Furthermore, these data were only produced in *Nicotiana benthamiana* for PVY-O and PVA and in *Physalis floridana* for PLRV. Similarly, for the virus transmission rates from aphid vector to host plant, there is only data available for the transmission of PVY, PVY-O, PVA, and PLRV by *Myzus persicae*. For PVY virus there is data on *Nicotiana benthamiana, N. tabacum*, and *Solanum tuberosum*; for PVY-O there is data on *N. benthamiana* and *S. tuberosum*; for PVA there is only data on *N. benthamiana*; and for PLRV there is data on *S. tuberosum* and *P. floridana*.

Aphids are the most common insects that efficiently spread viruses to crops. Of the 4400 know species, around 250 feeds on crops and of these approximately 200 have been identified as virus vectors and most of them are globally distributed. Aphids transmit around 275 virus species within 19 virus genera, representing more than 50% of insect-vectored viruses (Canto et al., 2009; Harrington et al., 2007). Aphids can transmit viruses in two main ways, non-circulative and circulative, due to their unique mouthpieces and other features (Ng and Perry, 2004). There is a great need to comprehensively fill the gaps on how temperature impacts life-history traits of aphids as well as the spread of viruses by them, in order to predict how global warming could impact aphid-borne diseases in crops. This knowledge could help to inform future pest and disease management strategies.

Changes in temperature may locally increase, decrease, or have no effect on the transmission of viruses by aphids. Mechanistic models allow us to assess the magnitude of these changes. It is critical to have data with enough replications to accurately estimate the optimum temperature and the thermal limits of virus transmission in crop ecosystems. Uncertainty in the models and in the predictions are driven by the amount of available data (Mordecai et al., 2019; Villena et al., 2022). Increases in temperatures may increase the geographical and seasonal ranges of aphid-borne diseases with high thermal optima and range of transmission (upper thermal minus the minimum thermal). However, increases in temperatures are more likely to shift or contract the geographic and seasonal ranges of aphid-borne diseases with lower thermal optima and small thermal range of transmission (Taylor et al., 2019; Villena et al., 2022). In our study, we appreciated that areas suitable for the transmission of PLRV by *A, gossypii* will shift to northern regions in all four warming scenarios, especially in Mexico, the north of Florida in the U.S., in Africa, and south-east Asia. The shift of PLRV transmission is greater for year 2100 under the SPP585 scenario. As mentioned before, there is need for data for other pathogens/vectors subsystems to build prediction models that could help to plan agriculture strategies and guaranty food for an increasing human population.

Our approach has some limitations, some of which could be addressed by extending these mechanistic models. One limitation is that in the estimation of the thermal optima and thermal limits for the life-history traits, virus transmission rates, and suitability models we used constant temperature data and did not incorporate the daily and seasonal temperature variations which are observed in nature. However, nonlinearities make it difficult to measure insect life-history traits and virus transmission rates, even at constant temperatures, especially at the thermal limits (Johnson et al., 2015; Mordecai et al., 2019; Villena et al., 2022). Another limitation is the non-inclusion of elevated concentration of CO_2_ which is an important compound for photosynthesis in plants. Carbon is a key element in the structure of plants. Increased CO_2_ concentrations enable faster growth due to carbon assimilation, although nutrient content could be diminished. Lower nutrient content could affect the feeding behavior of insects such as increasing sucking which could increase virus transmission. The atmospheric concentration of CO2 stood at 280 parts per million (ppm) during preindustrial times. However, in the present day, it has surged to nearly 420 ppm, with predictions suggesting it could climb as high as 1000 ppm under the RCP8.5 scenario (Nunes, 2023; Schwalm et al., 2020). Our trait-based thermal approach used to assess current state of aphids can be applied to understand the effects of global warming on many other types of vector-borne diseases in crops, including the ones transmitted by whiteflies, thrips, and psyllids.

The relationship between global warming, aphid populations, and aphid-mediated virus transmission to staple crops underscores the complex impact of climate change, especially global warming, on agriculture. A comprehensive understanding of these dynamics is essential for developing effective strategies to mitigate agricultural risks and ensure food security in the context of population growth, especially on the African and Asian continents. To achieve this goal, it is essential that more data be generated on the thermal responses of aphids and their transmission of pathogens. Through a combination of advanced research, innovative management practices, and proactive policy measures, the agriculture sector can adapt to the challenges posed by a warming world and safeguard crop production for future generations.

## Acknowledgments

This study was supported by the Earth Commons, Georgetown University’s Institute for Environmental and Sustainability. The authors thank the ECo Impact Awards for providing funding to aggregated existing data on the effects of temperature on aphid-borne disease dynamics on crops as well as for the generation of new data on life-history traits of *Myzus persicae* as well as the transmission rates of PVY from *M. persicae* to potatoes. The authors also want to thank the International Potato Center (CIP) and the International Centre for Insect Physiology and Ecology (*icipe*) for facilitating their laboratories to perform the above mentioned experiments.

