## Supplemental Material for "Thermal biology of aphids and implications for agriculture and food security"

**Section 1.** Thermal response of life-history traits and virus transmission rates

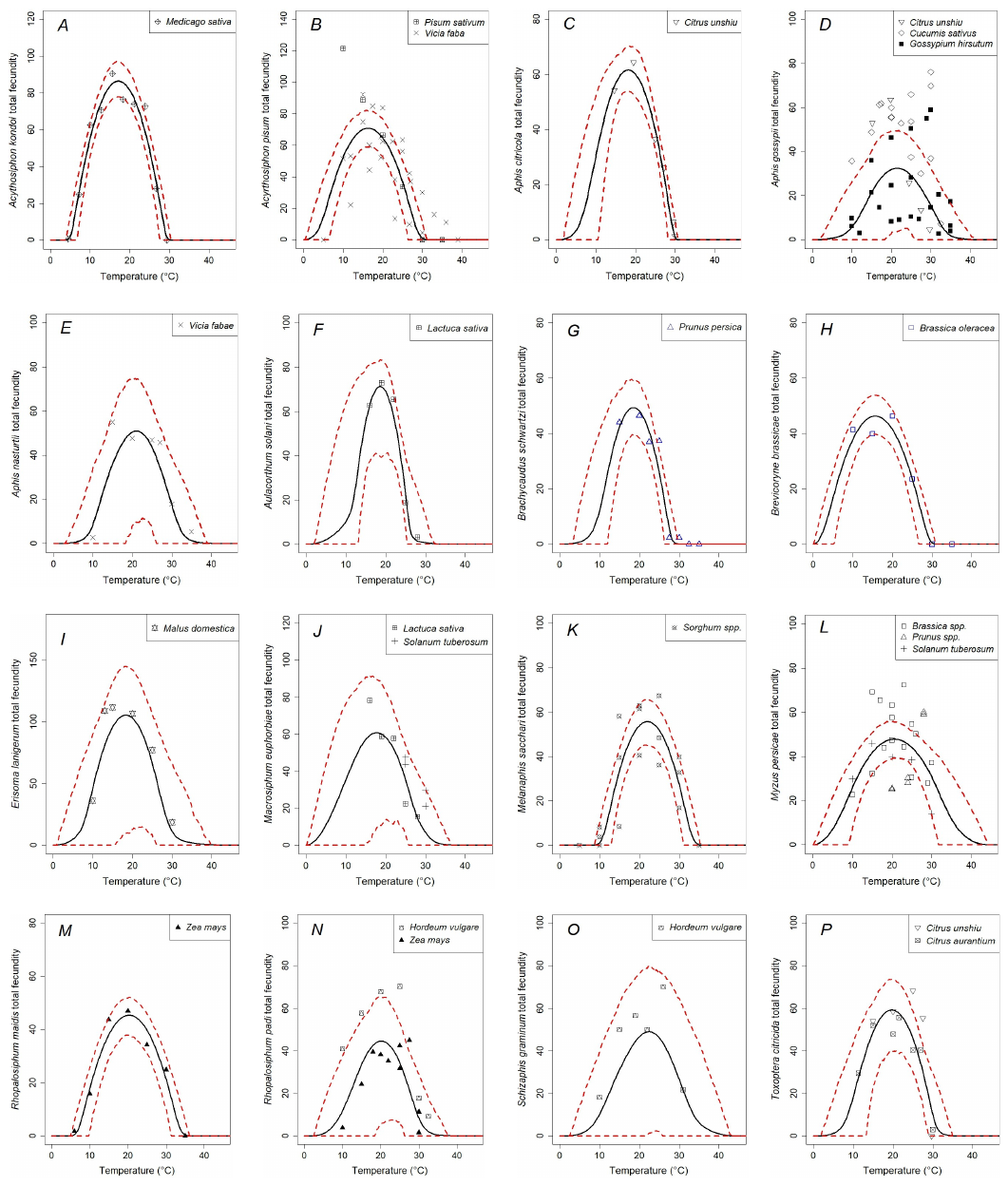

Fig. S1. Posterior mean (solid line) and 95% highest posterior density (HPD; dashed lines) of the thermal responses for aphids total fecundity (NFL) for: A) *Acyrthosiphom kondoi*, B) *Acyrtosiphon pisum*, C) *Aphis citricola*, D) *Aphis gossypii*, E) *Aphis nasturti*, F) *Aulacorthum solani*, G) *Brachycaudus schwartzi*, H) *Brevicoryne brassicae*, I) *Erisoma lanigerum*, J) *Macrosiphum euphorbiae*, K) *Melanaphis saccari*, L) *Myzus persicae*, M) *Rhopalosiphum maidis*, N) *Rhopalosiphum padi*, 0) *Schizaphis graminum*, and P) *Toxoptera citricida*.

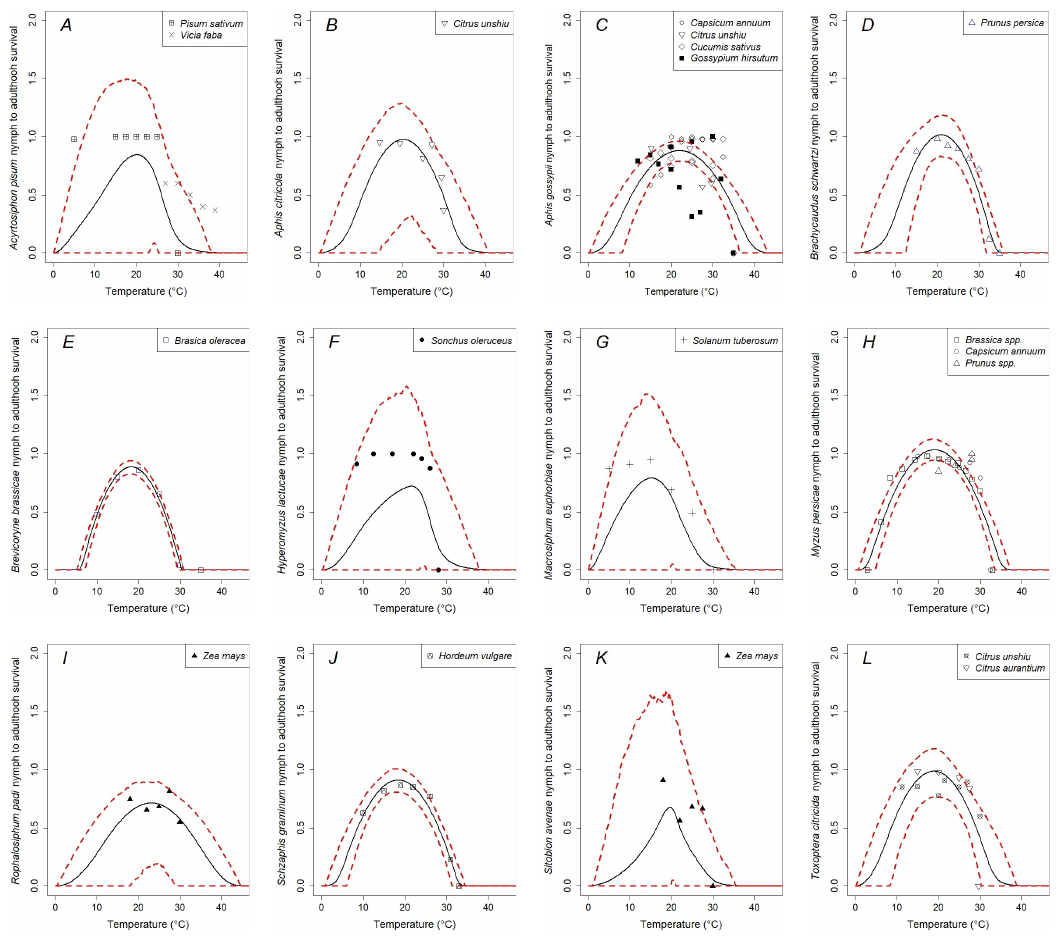

Fig. S2. Posterior mean (solid line) and 95% highest posterior density (HPD; dashed lines) of the thermal responses for the proportion of aphid nymphs surviving to adulthood (PNA) for: A) *Acyrthosiphon pisum* B) *Aphis citricola*, C) *Aphis gossypii*, D) *Brachycaudus schwartzi*, E) *Brevicoryne brassicae*, F) *Hyperomyzus lactucae*, G) *Macrosiphum euphorbiae*, H) *Myzus persicae*, I) *Rophalosiphum padi*, J) *Schizaphis graminum*, K) *Sitobion avenae*, and L) *Toxoptera citricida*.

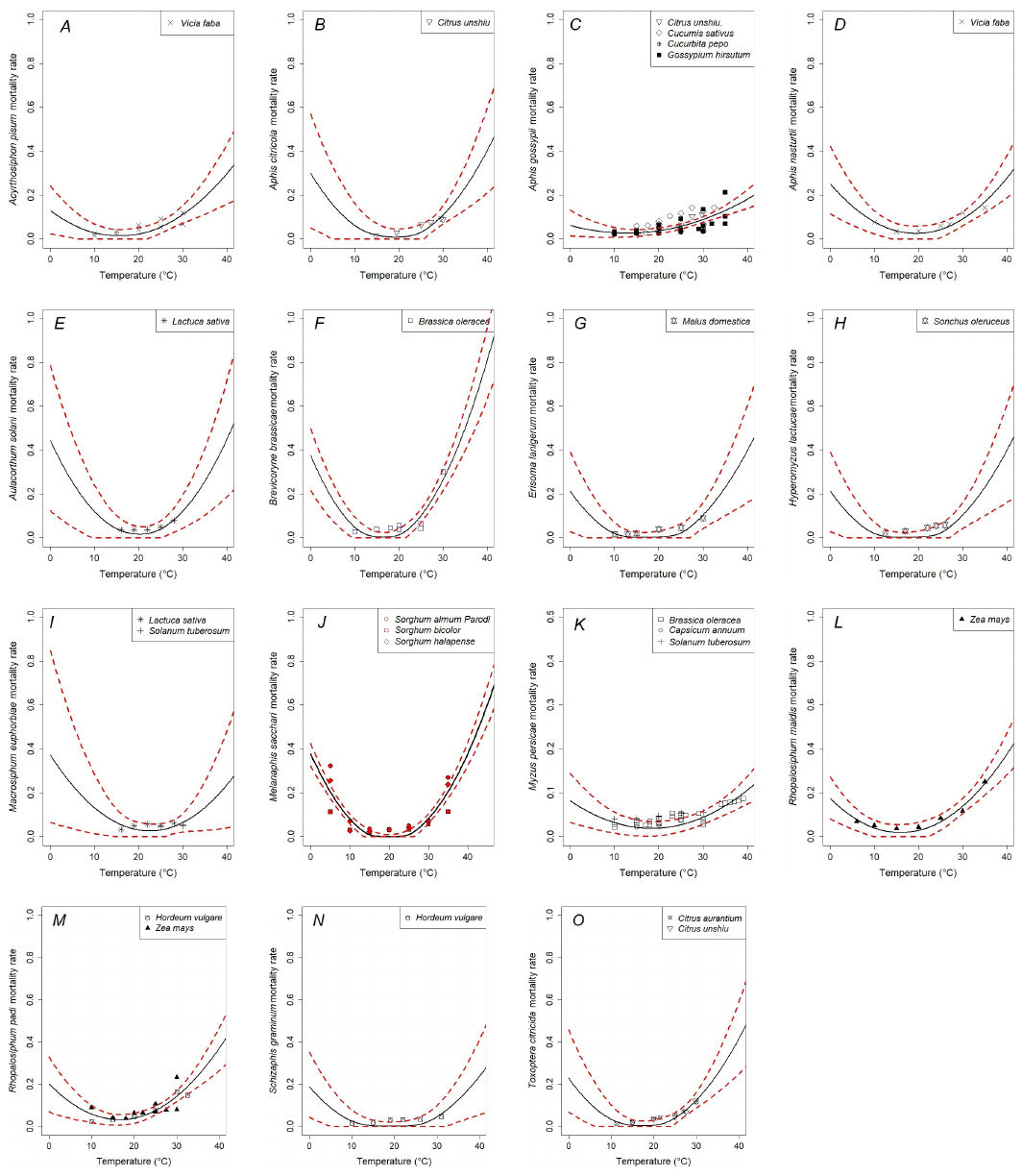

**Fig. S3.** Posterior mean (solid line) and 95% highest posterior density (HPD; dashed lines) of the thermal responses for aphid mortality rate (mu) for: A) *Acyrthosiphon pisum*, B) *Aphis citricola*, C) *Aphis gossypii*, D) *Aphis nasturtii*, E) *Aulacorthum solani*, F) *Brachycaudus schwartzi*, G) *Bravicoryne brassicae*, H) *Erisoma lanigerum*, I) *Hyperomyzus lactucae*, J) *Macrosiphum euphorbiae*, K) *Melanaphis sacchari*, L) *Myzus persicae*, M) *Rophalosiphum maidis*, N) *Rophalosiphum padi*, O) *Schizaphis graminum*, and P) *Toxoptera citricida*.

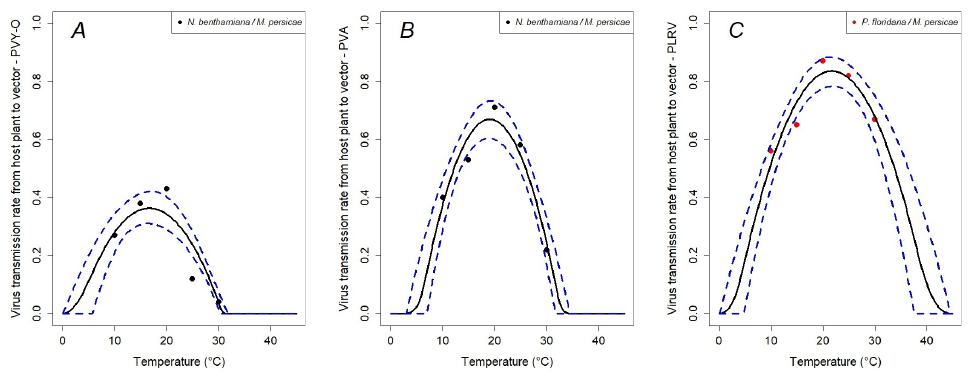

**Fig S4:** Posterior mean (solid line) and 95% highest posterior density (HPD; dashed lines) of the thermal responses for the transmission rates from host plant to *Myzus persicae* for the following viruses: A) potato virus Y strain O (PVY-O, B) potato virus A (PVA), and C) potato leaf roll virus (PLRV).

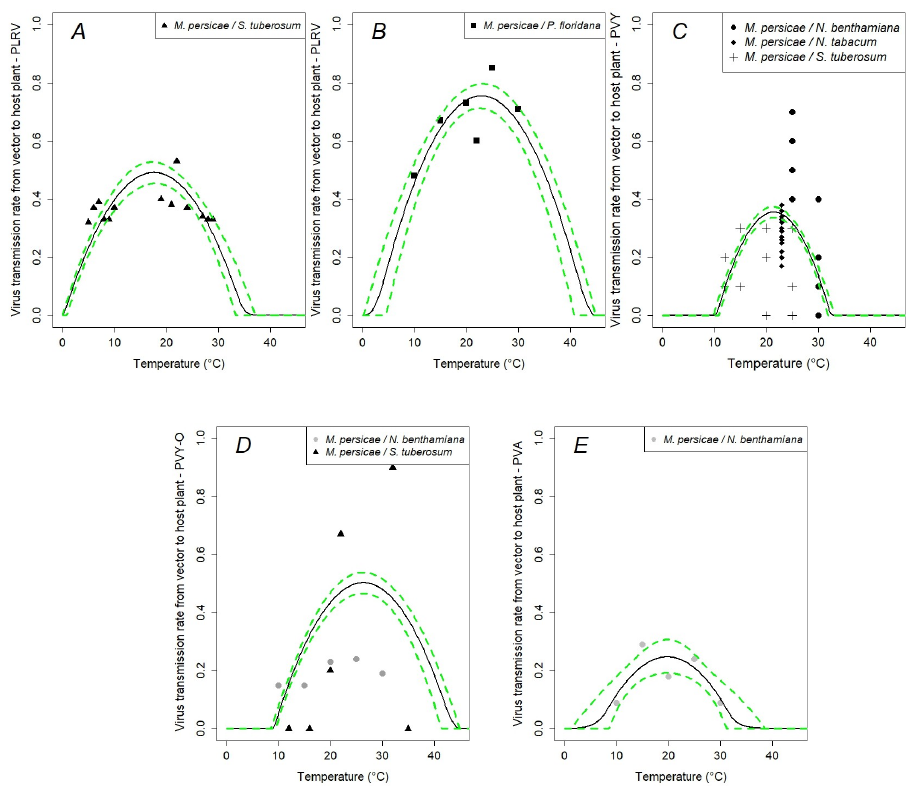

**FigS5:** Posterior mean (solid line) and 95% highest posterior density (HPD; dashed lines) of the thermal responses for the transmission rates from vector *M. persicae* to host plant for: A) potato leaf roll virus/*S. tuberosum*, B) potato leaf roll virus/*P. floridana*, C) potato virus Y, D) potato virus Y-O, and E) potato virus A.

**Section 2:** Optimum temperature and thermal limits for the different aphid species.

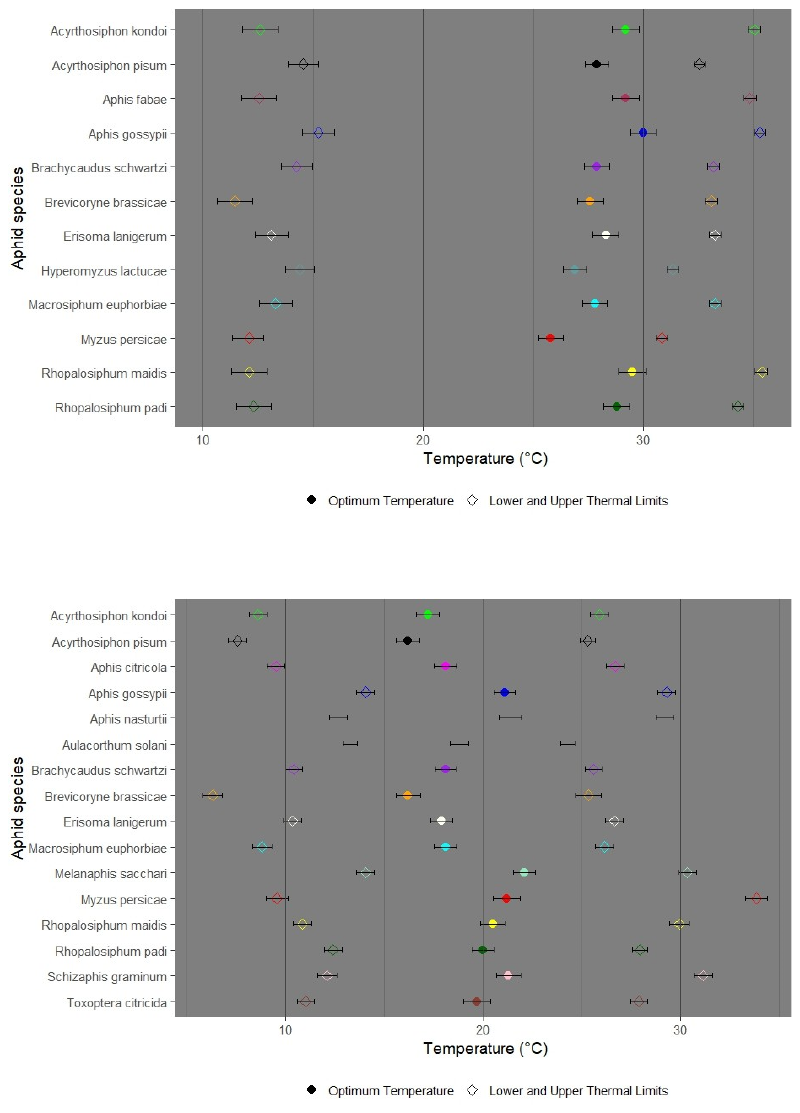

**Fig S6.** Posterior mean and credible intervals for the optimum temperature, and the lower and upper thermal responses for: Top panel: Aphid development rate (ADR); Bottom panel: Total aphid fecundity (NFL), expressed as the number of nymphs per female per life cycle. Symbols: optimum temperature: circles; thermal limits: diamonds, and 95% credible intervals: lines.

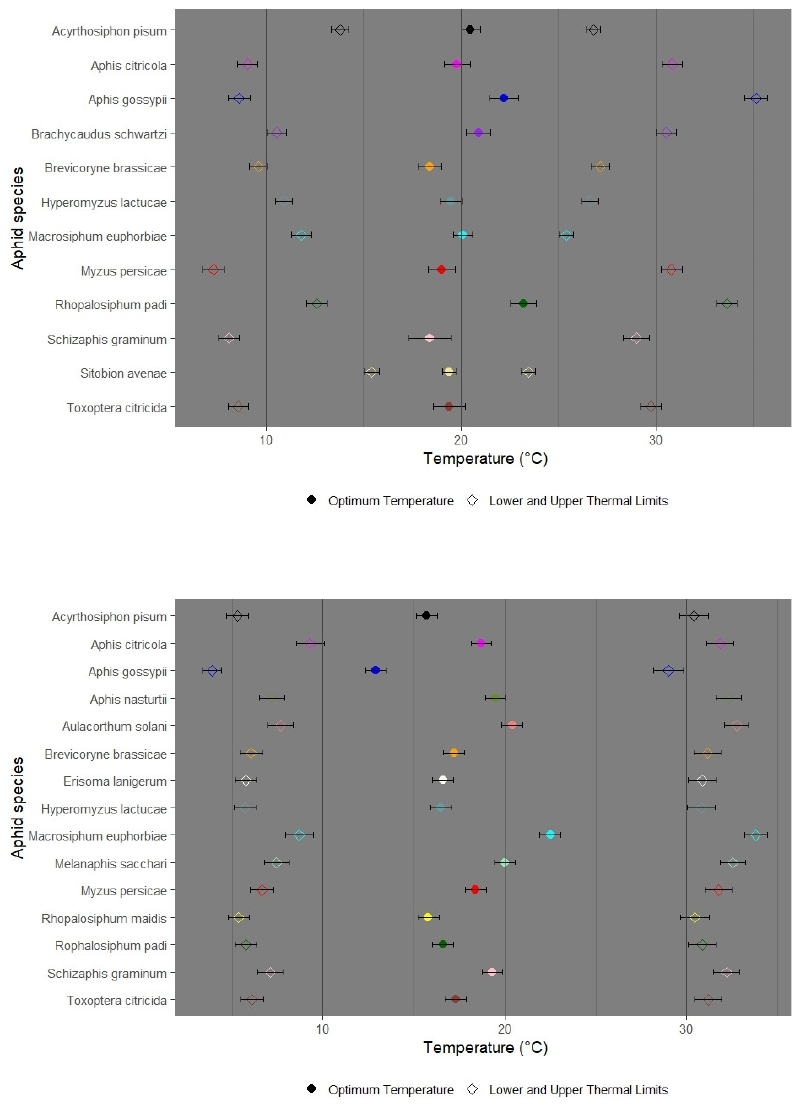

**Fig S7.** Posterior mean and credible intervals for the optimum temperature, and the lower and upper thermal responses for: Top panel: Proportion of nymphs reaching adulthood (PNA); Bottom panel: Mortality rate. Symbols: optimum temperature: circles; thermal limits: diamonds, and 95% credible intervals: lines.

**Section 3:** Map showing the global potato cropland.

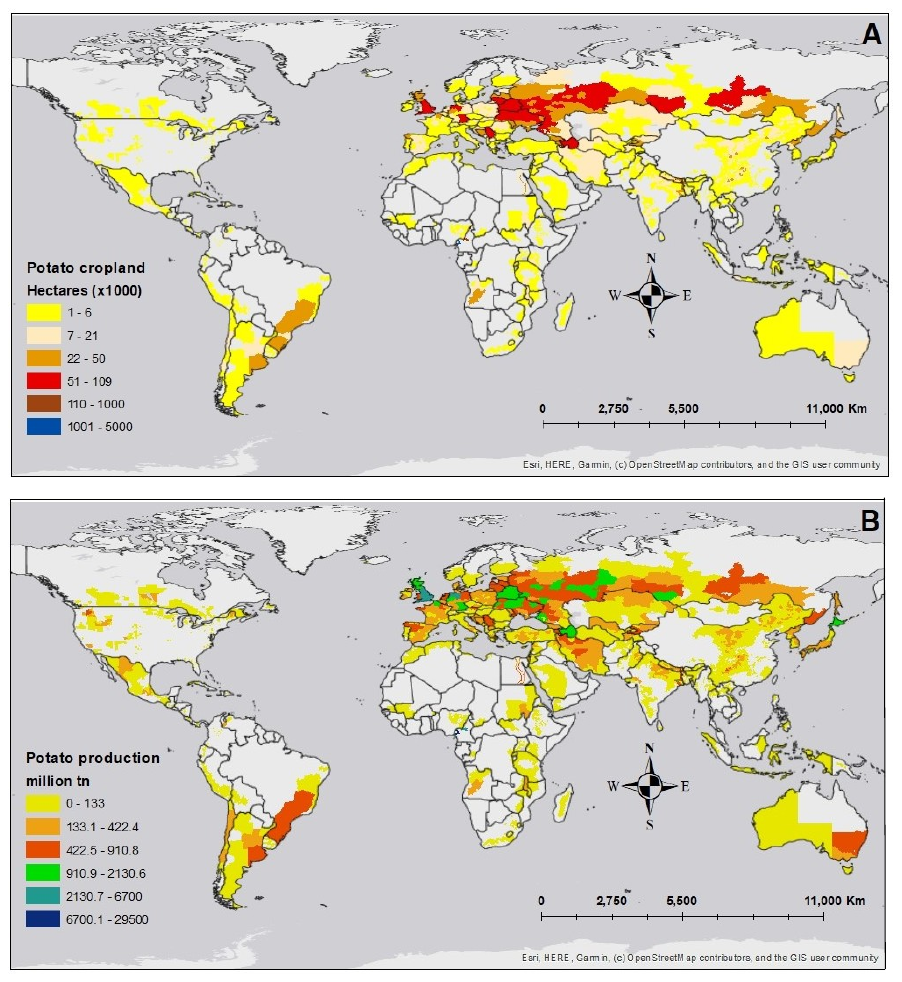

**Fig S8.** Global map showing A) potato cropland in thousands of hectares and B) potato production in million tons. These maps were elaborated by the International Potato Center (CIP).

**Section 4:** Year-round thermal suitability projections for the transmission of potato leaf roll virus (PLRV) by *Aphis gossypii*.

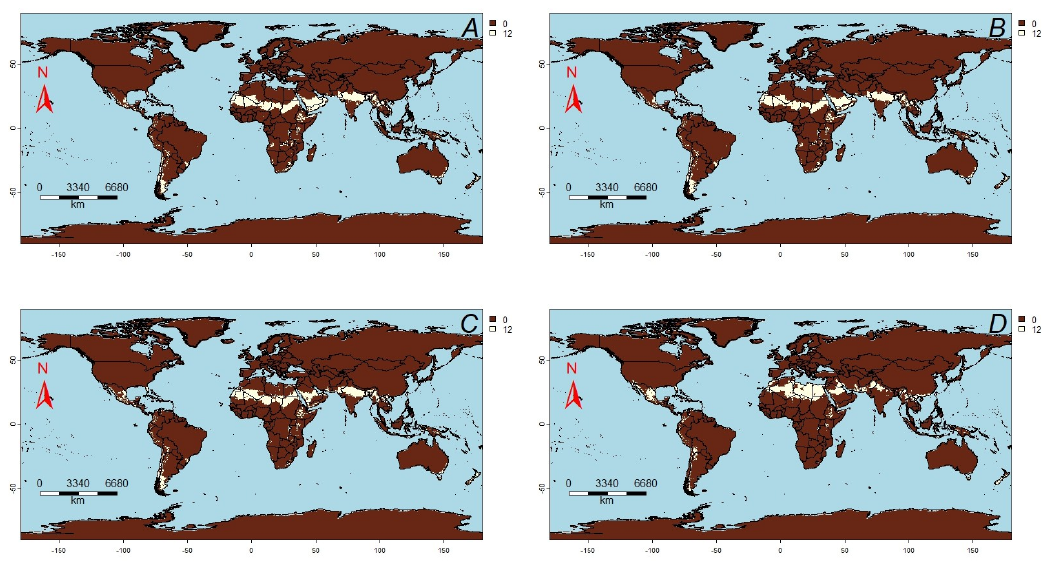

**Fig S9.** Year-round thermal suitability projection for the transmission of potato leaf roll virus (PLRV) by *Aphis gossypii* using the CMCC-ESM2 model from the Euro-mediterranean Centre on Climate Change. Year-round trans- mission suitability is shown for: A) year 2040 at SSP126 scenario, B) year 2040 at SSP585 scenario, C) year 2100 at SSP126 scenario, and D) year 2100 at SSP585 scenario. Year-round transmission is shown in white (12 months), where the posterior probability of S(T) *>* 0 is 0.95.

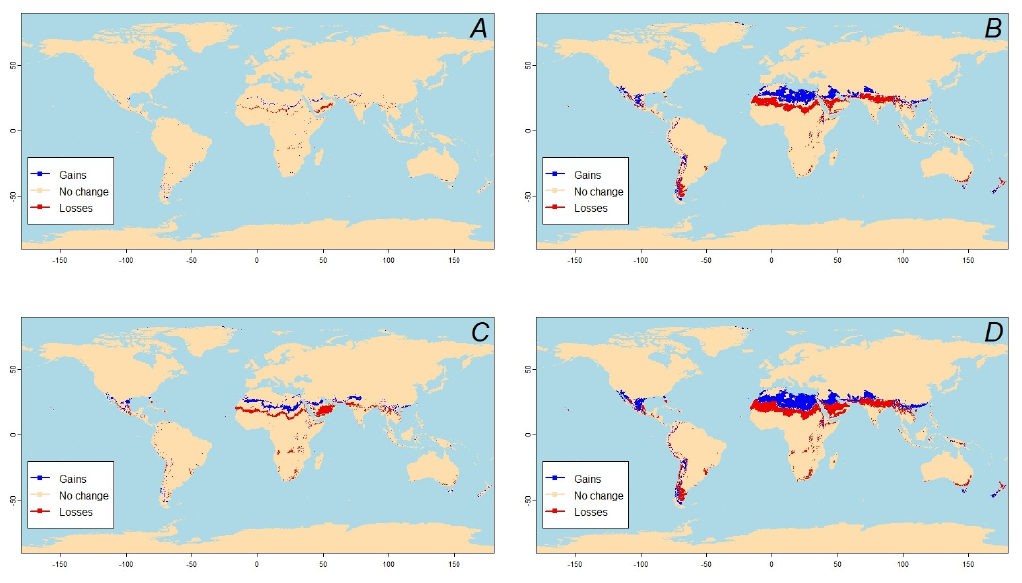

**Fig S10.** Changes in year-round thermal suitability projections for the transmission of potato leaf roll virus (PLRV) by *Aphis gossypii* using the CMCC-ESM2 model for: A) changes at year 2040 between SSP126 and SSP585 scenarios, B) changes at year 2100 between SSP126 and SSP585 scenarios, C) changes between years 2040 and 2100 for SSP126 scenario, and D) changes between years 2040 and 2100 for SSP585 scenario. Gains mean new areas where PLRV virus can be transmitted year-round by *Aphis gossypii*; Losses mean areas where PLRV virus cannot be transmitted year- round anymore by *Aphis gossypii*, where the posterior probability of S(T) *>* 0 is 0.95.

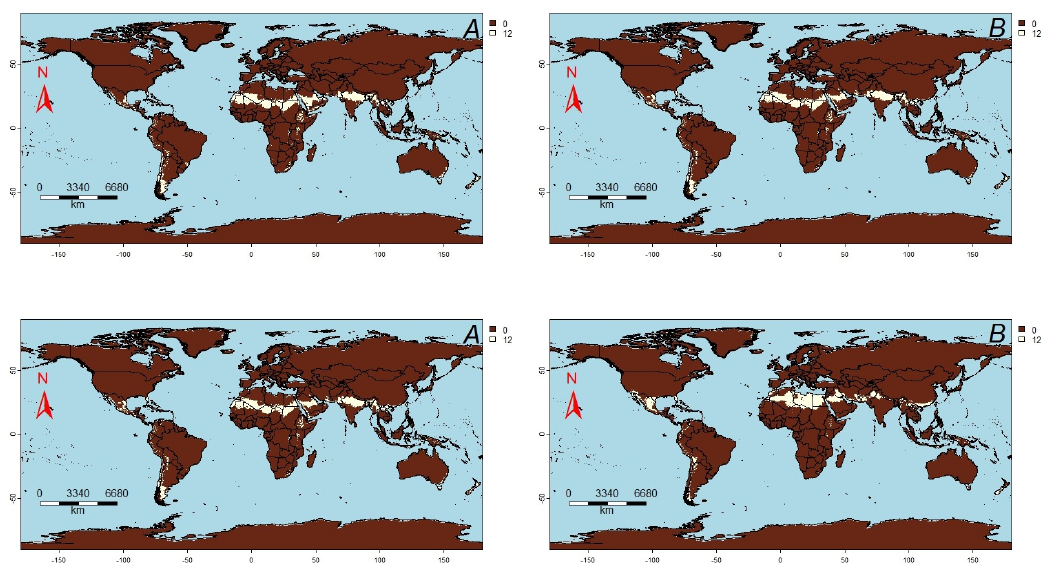

**Fig S11.** Year-round thermal suitability projection for the transmission of potato leaf roll virus (PLRV) by *Aphis gossypii* using the HadGEM3-GC31-LL model from the Hadley Centre for Climate Science and Services. Year-round transmission suitability is shown for: A) year 2040 at SSP126 scenario, B) year 2040 at SSP585 scenario, C) year 2100 at SSP126 scenario, and D) year 2100 at SSP585 scenario. Year-round transmission is shown in white (12 months), where the posterior probability of S(T) *>* 0 is 0.95.

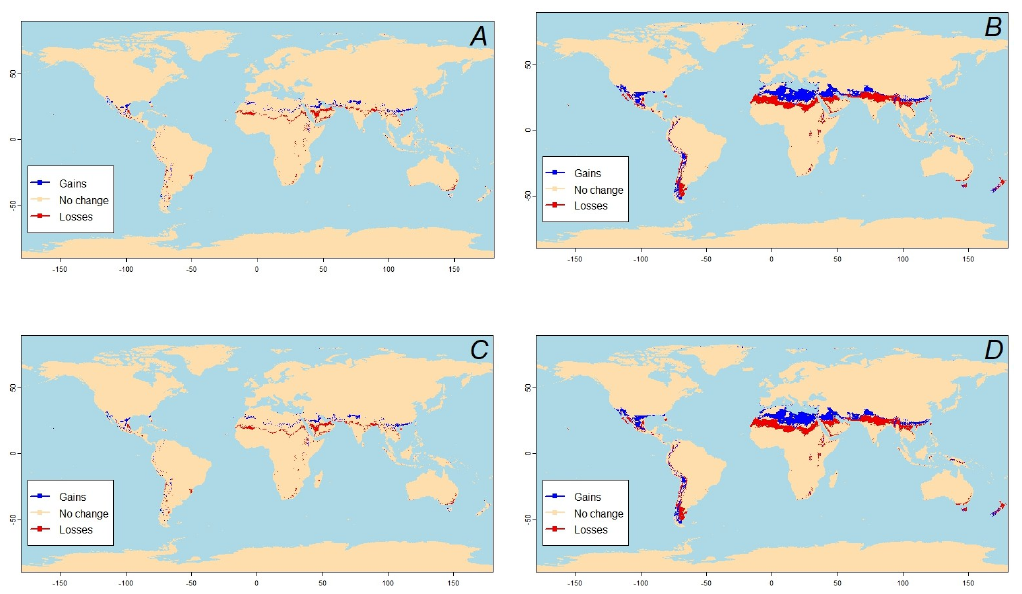

**Fig S12.** Changes in year-round thermal suitability projections for the transmission of potato leaf roll virus (PLRV) by *Aphis gossypii* using the HadGEM3-GC31-LL model for: A) changes at year 2040 between SSP126 and SSP585 scenarios, B) changes at year 2100 between SSP126 and SSP585 scenarios, C) changes between years 2040 and 2100 for SSP126 scenario, and D) changes between years 2040 and 2100 for SSP585 scenario. Gains mean new areas where PLRV virus can be transmitted year-round by *Aphis gossypii*; Losses mean areas where PLRV virus cannot be transmitted year-round anymore by *Aphis gossypii*, where the posterior probability of S(T) *>* 0 is 0.95.

**Section 5:** Map showing the locations where the different studies took place.

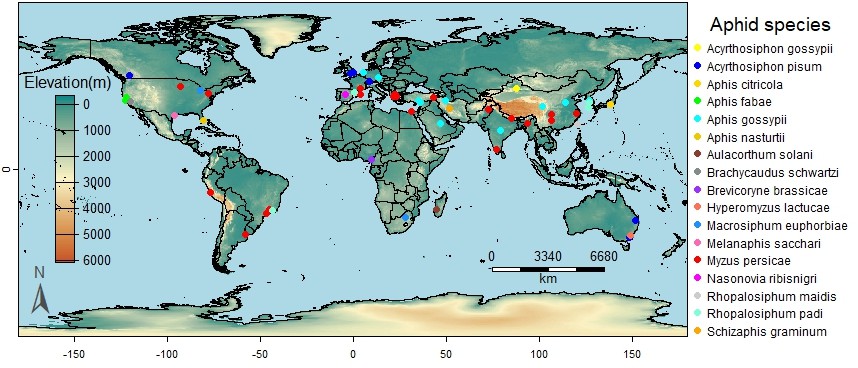

**Fig S13.** Map showing the locations where the different studies that produced life-history traits for 17 aphid species took place.

**Section 6:** Tables summarizing the optimum temperature and the thermal limits for the different life-history traits

Table S1. Average thermal optima, thermal limits, and 95% confidence intervals (Cis) for aphid fecundity.

| **Species** | **Crops** | **Lower thermal (CIs)** | **Optimum Temp**  **(CIs)** | **Upper thermal**  **(CIs)** |
| --- | --- | --- | --- | --- |
| *Acyrthosiphon kondoi* | *Medicago sativa* | 8.6 | 17.2 | 25.9 |
|  |  | (8.14, 9.06) | (16.61, 17.79) | (25.44, 26.36) |
| *Acyrthosiphon pisum* | *Pisum sativum*, | 7.55 | 16.2 | 25.3 |
|  | *Vicia faba* | (7.08, 8.02) | (15.6, 16.8) | (24.93, 25.67) |
| *Aphis citricola* | *Citrus unshiu* | 9.5 | 18.1 | 26.7 |
|  |  | (9.04, 9.96) | (17.51, 18.69) | (26.24, 27.16) |
|  | *Citrus unshiu*, |  |  |  |
| *Aphis gossypii* | *Cucumis sativus*, *Gossypium hirsutum* | 14.05  (13.6, 14.5) | 21.1  (20.56, 21.64) | 29.3  (28.84, 29.76) |
| *Aphis nasturtii* | *Vicia faba* | 12.65 | 21.4 | 29.2 |
|  |  | (12.2, 13.1) | (20.82, 21.98) | (28.75, 29.65) |
| *Brachycaudus schwartzi* | *Prunus persica* | 10.4 | 18.1 | 25.6 |
|  |  | (9.97, 10.83) | (20.85, 21.95) | (25.17, 26.03) |
| *Aulacorthum solani* | *Lactuca sativa* | 13.25 | 18.8 | 24.3 |
|  |  | (12.88, 13.62) | (18.33, 19.27) | (23.93, 24.67) |
| *Brevicoryne brassicae* | *Brassica oleracea* | 6.3 | 16.2 | 25.35 |
|  |  | (5.81, 6.79) | (15.58, 16.82) | (24.72, 25.98) |
| *Erisoma lanigerum* | *Malus domestica* | 10.35 | 17.9 | 26.65 |
|  |  | (9.91, 10.79) | (17.33, 18.47) | (26.19, 27.11) |
| *Macrosiphum euphorbiae* | *Lactuca sativa*, | 8.8 | 18.1 | 26.15 |
|  | *Solanum tuberosum* | (8.31, 9.29) | (17.53, 18.67) | (25.68, 26.62) |
|  | *Brassica oleracea*, |  |  |  |
| *Myzus persicae* | *Prunus armeniaca*, | 9.55 | 21.2 | 33.85 |
|  | *Prunus persica* |  |  |  |
|  |  | (9, 10.1) | (20.51, 21.89) | (33.3, 34.4) |
| *Rhopalosiphum maidis* | *Zea mays* | 10.85 | 20.5 | 29.95 |
|  |  | (10.37, 11.33) | (19.88, 21.12) | (29.47, 30.43) |
|  | *Sorghum bicolor*, |  |  |  |
| *Melanaphis sacchari* | *Sorghum halepense*, *Sorghum ssp.* | 14.05  (13.6, 14.5) | 22.1  (21.53, 22.67) | 30.35  (29.9, 30.8) |
| *Schizaphis graminum* | *Hordeum vulgare* | 12.1 | 21.3 | 31.15 |
|  |  | (11.61, 12.59) | (20.68, 21.92) | (30.67, 31.63) |
| *Toxoptera citricida* | *Citrus aurantium*, | 11 | 19.7 | 27.9 |
|  | *Citrus unshiu* | (10.57, 11.43) | (19, 20.4) | (27.48, 28.32) |
| *Rhopalosiphum padi* | *Hordeum vulgare*,  *Zea mays* | 12.4  (11.95, 12.85) | 20  (19.45, 20.55) | 27.95  (27.56, 28.34) |

Table S2. Average thermal optima, thermal limits, and 95% confidence intervals (CIs) for the proportion of aphid nymphs that reach adulthood

| **Species** | **Crops** | **Lower thermal**  **(CIs)** | **Optimum Temp**  **(CIs)** | **Upper thermal**  **(CIs)** |
| --- | --- | --- | --- | --- |
| *Acyrthosiphon pisum* | *Pisum sativum,*  *Vicia faba* | 13.8  (13.35, 14.25) | 20.5  (19.99, 21.01) | 26.8  (26.45, 27.15) |
| *Aphis citricola* | *Citrus unshiu* | 9.05 | 19.8 | 30.85 |
|  |  | (8.52, 9.58) | (19.14, 20.46) | (30.33, 31.37) |
|  | *Capsicum annuum*, |  |  |  |
|  | *Citrus unshiu*, |  |  |  |
| *Aphis gossypii* | *Cucumis sativus*, *Gossypium hirsutum* | 8.6  (8.03, 9.17) | 22.2  (21.47, 22.93) | 35.15  (34.58, 35.72) |
| *Brachycaudus schwartzi* | *Prunus persica* | 10.55 | 20.9 | 30.55 |
|  |  | (10.05, 11.05) | (20.27, 21.53) | (30.05, 31.05) |
| *Brevicoryne brassicae* | *Brassica oleracea* | 9.6 | 18.4 | 27.15 |
|  |  | (9.14, 10.06) | (17.08, 19) | (26.68, 27.62) |
| *Hyperomyzus lactucae* | *Sonchus oleruceus* | 10.9 | 19.5 | 26.65 |
|  |  | (10.47, 11.33) | (18.94, 20.06) | (26.22, 27.08) |
| *Macrosiphum euphorbiae* | *Solanum tuberosum* | 11.8 | 20.1 | 25.4 |
|  |  | (11.28, 12.32) | (19.61, 20.59) | (25.02, 25.78) |
|  | *Brassica spp.*, |  |  |  |
| *Myzus persicae* | *Capsicum annuum*, *Prunus spp.* | 7.3  (6.76, 7.84) | 19  (18.31, 19.69) | 30.8  (30.26, 31.34) |
| *Rhopalosiphum padi* | *Zea mays* | 12.6 | 23.2 | 33.65 |
|  |  | (12.09, 13.11) | (22.55, 23.85) | (33.15, 34.15) |
| *Schizaphis graminum* | *Hordeum vulgare* | 8.1 | 18.4 | 29 |
|  |  | (7.59, 8.61) | (17.33, 19.47) | (28.33, 29.67) |
| *Sitobion avenae* | *Zea mays* | 15.4 | 19.4 | 23.45 |
|  |  | (15.01, 15.79) | (19.05, 19.75) | (23.1, 23.8) |
| *Toxoptera citricida* | *Citrus aurantium*, | 8.55 | 19.4 | 29.75 |
|  | *Citrus unshiu* | (8.03, 9.07) | (18.56, 20.24) | (29.24, 30.26) |

**Table S3.** Average thermal optima, thermal limits, and 95% confidence intervals (CIs) for aphid mortality rate

| **Species** | **Crops** | **Lower thermal**  **(CIs)** | **Optimum Temp**  **(CIs)** | **Upper thermal**  **(CIs)** |
| --- | --- | --- | --- | --- |
| *Acyrthosiphon pisum* | *Vicia faba* | 5.3  (4.71, 5.89) | 15.7  (15.13, 16.27) | 30.4  (29.61, 31.18) |
| *Aphis citricola* | *Citrus unshiu* | 9.3 | 18.7 | 31.85 |
|  |  | (8.52, 10.07) | (18.13, 19.27) | (31.12, 32.58) |
|  | *Citrus unshiu*, |  |  |  |
|  | *Cucumis sativus*, |  |  |  |
| *Aphis gossypii* | *Cucurbita pepo*,  *Gossypium hirsutum* | 3.9  (3.4, 4.4) | 12.9  (12.33, 13.47) | 29  (28.16, 29.84) |
| *Aphis nasturtii* | *Vicia faba* | 7.2 | 18.93 | 32.3 |
|  |  | (6.52, 7.88) | (19.83, 20.07) | (31.6, 33) |
| *Aulacorthum solani* | *Lactuca sativa* | 7.65  (6.95, 8.35) | 20.4  (19.83, 20.97) | 32.75  (32.07, 33.43) |
| *Brevicoryne brassicae* | *Brasica oleracea* | 6.05  (5.42, 6.67) | 17.2  (16.63, 17.77) | 30.39  (30.39, 31.9) |
| *Erysoma lanigerum* | *malus domestica* | 5.75  (5.14, 6.36) | 16.6  (16.03, 17.17) | 30.85  (30.08, 31.62) |
| *Hyperomyzus lactucae* | *Sonchus oleruceus* | 5.7 | 16.5 | 30.8 |
|  |  | (5.09, 6.31) | (15.93, 17.07) | (30.03, 31.57) |
| *Macrosiphum euphorbiae* | *Solanum tuberosum* | 8.7 | 22.5 | 33.8 |
|  |  | (7.95, 9.45) | (21.93, 23.07) | (33.17, 34.43) |
|  | *sorghum almum*, |  |  |  |
| *Melanaphis sacchari* | *sorghum bicolor*, *sorghum halapense* | 7.45  (6.76, 8.14) | 20  (19.43, 20.57) | 32.55  (31.86, 33.24) |
|  | *Brassica oleracea*, |  |  |  |
| *Myzus persicae* | *Capsicum annuum*,  *Solanum tuberosum* | 6.65  (5.99, 7.3) | 18.4  (17.83, 18.97) | 31.75  (31.02, 32.48) |
| *Rhopalosiphum maidis* | *Zea mays* | 5.35 | 15.8 | 30.45 |
|  |  | (4.76, 5.35) | (15.23, 16.37) | (29.67, 31.23) |
| *Rhopalosiphum padi* | *Hordeum vulgare*,  *Zea mays* | 5.75  (5.14, 6.36) | 16.6  (16.03, 17.17) | 30.85  (30.08, 31.62) |
| *Schizaphis graminum* | *Hordeum vulgare* | 7.1 | 19.3 | 32.2 |
|  |  | (6.42, 7.78) | (18.73, 19.87) | (31.49, 32.91) |
| *Toxoptera citricida* | *Citrus aurantium*, | 6.1 | 17.3 | 31.2 |
|  | *Citrus unshiu* | (5.47, 6.73) | (16.73, 17.87) | (30.45, 31.95) |

Section 7. Available data on life-history traits of aphids that transmit pathogens to crops.

| **Scientific Name(s)** | **Traits** | **Crop Species** | **Pathogen** | **Reference** |
| --- | --- | --- | --- | --- |
| *Acyrthosiphon kondoi* | adt, adr, nfl | *Medicago sativa Cv. Moapa 69* |  | Summers et al. 1984. |
| *Acyrthosiphon pisum* | adt, adr, lgv, mu, nfl | *Vicia faba* |  | Ahn et al. 2020. |
| *Acyrthosiphon pisum* | adt | *Medicago sativa* |  | Campbell and MacKauer, 1975. |
| *Acyrthosiphon pisum* | adt |  |  | Lamb and MacKay, 1998. |
| *Acyrthosiphon pisum* | adt, lgv, nfl | *Pisum sativum* |  | Lu and Kuo, 2008. |
| *Acyrthosiphon pisum* | adt, lgv, mu, nfl | *Vicia faba* |  | Mastoi et al. 2020. |
| *Acyrthosiphon pisum* | adt, nfl | *Vicia faba* |  | Morgan et al. 2001. |
| *Acyrthosiphon pisum* | adt, lgv, mu, nfl | *Pisum sativum* |  | Siddiqui et al. 1973. |
| *Acyrthosiphon pisum* | adt | *Vicia faba* |  | Mala et al. 2023. |
| *Acyrthosiphon pisum* | adt, adr, nfl | *Pisum sativum* |  | Bieri et al. 1983. |
| *Acyrthosiphon gossypii* | lgv, mu, nfd, nfl | *Gossypium hirsutum L. cv. Zhongmian* |  | Liu et al. 2021. |
| *Aphis citricola* | lgv, mu, nfl, pna | *Citrus unshiu* |  | Komasaki, S. 1982. |
| *Aphis fabae* | lgv, nfl | *Brassica kaber Malva parviflora Veronica hederifolia Convolvulus arvensis Solanum nigrum Amaranthus retroflexus Capsella bursa-pastoris Amsinckia intermedia Chenopodium album* |  | Fernandez-Quintanilla et al. 2002. |
| *Aphis fabae* | adt, adr, nfl | *Vicia faba* |  | Tsitsipis and Mittler, 1976. |
| *Aphis gossypii* | lgv, mu, nfl, pna | *Citrus unshiu* |  | Komasaki, S. 1982. |
| *Aphis gossypii* | adt, adr, lgv, mu | *Cucurbita pepo* |  | Aldyhim and Khalil. 1993. |
| *Aphis gossypii* | adt, adr, lgv, mu, nfd, pna | *Gossypium hirsutum* |  | Kersting et al. 1999. |
| *Aphis gossypii* | adt, adr, lgv, mu, nfl | *Cucumis sativus* |  | Kocourek et al. 1994. |
| *Aphis gossypii* | adt, adr, mu, pna | *Capsicum annuum* |  | Satar et al. 2008. |
| *Aphis gossypii* | adt, adr, nfd, pna | *Cucumis sativus* |  | Satar et al. 2005. |
| *Aphis gossypii* | adt, adr, lgv, mu, nfd, nfl, pna | *Cucumis sativus* |  | Steenis and El-Khawass. 1995. |
| *Aphis gossypii* | adt, adr, dt, lgv, mu, nfd, nfl | *Gossypium hirsutum* |  | Xia et al. 1999. |
| *Aphis gossypii* | adt, adr, dt, lgv, mu, pna | *Cucumis sativus* |  | Zamani et al. 2006. |
| *Aphis gossypii* | adt, adr, nfl | *Hybiscus syriacus* |  | Hosseini-Tabesh et al. 2015. |
| *Aphis gossypii* | adt, lgv, mu, nfd, nfl | *Cucumis sativus* |  | Kim et al. 2004. |
| *Aphis gossypii* | Lgv, mu, nfd, nfl | *Gossypium hirsutum L. cv. Zhongmian* |  | Liu et al. 2021. |
| *Aphis gossypii* | adt, adr | *Cucumis sativus cv. Negin* |  | Zamani et al. 2007. |
| *Aphis gossypii* | adr, adt, nfl, pna | *Gossypium hirsutum* |  | Nagrare et al. 2021. |
| *Aphis gossypii* | adt, lgv, mu, nfl | *Gossypium hirsutum L. cv. Paymaster 2326RR* |  | Parajulee, 2007. |
| *Aphis gossypii* | adt, adr | *Cucumis sativus* |  | Zamani et al. 2007. |
| *Aphis gossypii* | trh | *Solanum tuberosum L. cv. BSS-341,*  *Capsicum annuum L. cv. Demre,*  *Lycopersicon esculentum cv. H-2275,*  *Physalis floridana* | PVY, PLRV | Sertkaya and Sertkaya, 2005. |
| *Aphis nasturtii* | adt, lgv, nfl | *Vicia faba* |  | Wang et al. 1997. |
| *Aulacorthum solani* | adt, gt, lgv, nfl | *Lactuca sativa* |  | Conti et al. 2009. |
| *Brachycaudus schwartzi* | adt, adr, lgv, mu, nfl, pna | *Prunus persica* |  | Satar et al. 2002. |
| *Brevicoryne brassicae* | lgv | *Brassica oleracea* |  | Gupta et al. 2022. |
| *Brevicoryne brassicae* | adt, adr, lgv, mu, nfl, pna | *Brassica oleracea var Marcanta* |  | Soh et al. 2018. |
| *Erisoma lanigerum* |  |  |  |  |
| *Hyperomyzus lactucae* | adt, adr, lgv, mu, pna | *Sonchus oleraceus* |  | Shu-sheng and Hughes, 1987. |
| *Hyperomyzus lactucae* | adt, adr, lgv, mu, pna | *Ribes nigrum* | SYVV, LNYV | Shu-sheng and Hughes, 1987. |
| *Macrosiphum euphorbiae* | adt, lgv, nfd, nfl | *Solanum tuberosum* |  | Beetge & Krüger, 2019. |
| *Macrosiphum euphorbiae* | lgv, nfl | *Lactuca sativa* |  | Conti et al. 2009. |
| *Macrosiphum euphorbiae* | adt, mu | *Nicotiana tabacum L., Solanum tuberosum* |  | Barlow, 1962. |
| *Macrosiphum rosae (L.)* | adt, lgv, nfl | *Rosa rubiginosa* |  | Olmez et al. 2003. |
| *Melanaphis sacchari* | lgv, nfd, nfl | *Sorghum bicolor cv. KS 585* |  | Souza et al. 2019. |
| *Myzus persicae* | adt, ndf, lgv |  |  | Asante et al. 1991. |
| *Myzus persicae* | adr, nfd | *Brassica rapa,*  *Solanum tuberosum* |  | Davis et al. 2006. |
| *Myzus persicae* | adt, adr, mu, pna | *Capsicum annuum* |  | Satar et al. 2008. |
| *Myzus persicae* | adt, adr, lgv, mu, nfd, nfl | *Brassica oleracea var. Capitata* |  | Baral et al. 2021. |
| *Myzus persicae* | adt, adr, lgv, mu, nfl | *Brassica oleracea* |  | Chanu and Gupta, 2014. |
| *Myzus persicae* | adt, adr | *Capsicum annuum* |  | Zamani et al. 2007. |
| *Myzus persicae* | trh, trv | *Nicotiana benthamiana, Physalis floridana* | PVY-O, PVA, PLRV | Chung et al. 2016. |
| *Myzus persicae* | Lgv, mu, nfd | *Capsicum annuum cv Ligia* |  | Rodrigues et al. 2010. |
| *Myzus persicae* | Lgv, mu, nfd, nfl | *Brassica oleracea var acephala* |  | Cividanes et al. 2003. |
| *Myzus persicae* | adt, adr, mu, pna | *Brassica campestris* |  | Liu & Meng, 1999. |
| *Myzus persicae* | nfd | *Capsicum annuum* | PCV-1 | Safari et al. 2019. |
| *Myzus persicae* | trh | *Solanum tuberosum L.* | PLRV | Ifitkhar et al. 2020. |
| *Myzus persicae* | adt, lgv, nfl | *Vicia faba Capsicum annuum Raphanus sativus Nicotiana tabacum Brassica campestris* |  | Hong et al. 2019. |
| *Myzus persicae* | adt, lgv | *Capsicum annuum (pepper)* |  | La Rossa, 2013. |
| *Myzus persicae* | adt, lgv, nfl | *Brassica napus L. Brassica oleracea L. var. acephala DC Raphanus sativus L. Brassica pekinensis L. Nicotiana tabacum L.* |  | Jiang et al. 2022. |
| *Myzus persicae* | adr, adt, lgv, nfl | *Capsicum annuum* |  | Özgökçe et al. 2018. |
| *Myzus persicae* | lgv | *Brassica oleracea* |  | Gupta et al. 2022. |
| *Myzus persicae* | trh | *Solanum tuberosum L. cv. BSS-341,*  *Capsicum annuum L. cv. Demre,*  *Lycopersicon esculentum cv. H-2275,*  *Physalis floridana* | PVY, PLRV | Sertkaya and Sertkaya, 2005. |
| *Myzus persicae* | adt, adr | *Capsicum annuum* |  | Komasaki, S. 1982. |
| *Myzus persicae* | adt, lgv | *Brassica oleracea* |  | Saleesha et al. 2022. |
| *Myzus persicae* | adt, mu | *Nicotiana tabacum L., Solanum tuberosum* |  | Barlow, 1962. |
| *Myzus persicae* | adr, adt, nfd, nfl, pna | *Prunus armeniaca,*  *Prunus persica* |  | Abdel-Salam et al. 2009. |
| *Myzus persicae* | lgv, nfl | *Brassica kaber, Malva parviflora, Veronica hederifolia, Convolvulus arvensis, Solanum nigrum, Amaranthus retroflexus, Capsella bursa-pastoris, Amsinckia intermedia, Chenopodium album* |  | Fernandez-Quintanilla et al. 2002. |
| *Myzus persicae* | adt, lgv, nfl | *Capsicum annuum* |  | Özgökçe et al. 2023. |
| *Myzus persicae* | trh | *Nicotiana benthamiana* | PVY | del Toro et al. 2019. |
| *Myzus persicae* | trh | *Beta vulgaris,*  *Brassica napus* | BMYV, TuYV | Armand et al. 2021 |
| *Myzus persicae* | adt, lgv, nfl | *Nicotiana tabacum* |  | Goundoudaki et al. 2003. |
| *Myzus persicae* | adt, lgv, mu, nfl | *Nicotiana tabacum L., Capsicum annum L.* |  | Nikolakakis et al. 2003. |
| *Myzus persicae* | trh | *Nicotiana tabacum* | PVY | Kanavaki et al. 2006. |
| *Nasonovia ribisnigri* | adt | *Lactuca sativa var. longifolia* |  | Maria Diaz et al. 2005. |
| *Rhopalosiphum padi* | adt, dt, lgv, nfd, nfl | *Brachiaria ruziziensis* |  | Auad et al., 2009. |
| *Rhopalosiphum padi* | adt, lgv, nfl | *Zea mays L.* |  | Kuo, 2006. |
| *Rhopalosiphum padi* | lgv, mu, nfl | *Zea mays cv. Prisma G-4730* |  | Asin and Pons, 2001. |
| *Rhopalosiphum padi* | adr, adt, lgv, nfd, nfl | *Hordeum vulgare var. Hyemi* |  | Park et al. 2017. |
| *Rhopalosiphum padi* | adt, adr, lgv, nfd, nfl |  |  | Park et al. 2016. |
| *Schizaphis graminum* | adt, lgv, nfd, nfl | *Hordeum vulgare L.* |  | Tofangsazi et al. 2012. |
| *Toxoptera citricida* | lgv, mu, nfl, pna | *Citrus unshiu* |  | Zamani et al. 2007. |
| *Tuberolachnus salignus* | adr, lgv, nfd, nfl | *Salix viminalis “Q683”* |  | Matilda and Leather, 2001. |
| *Uroleucon ambrosiae* | adt, lgv, nfl | *Lactuca sativa* |  | Conti et al. 2009. |

<https://doi.org/10.1079/BER200062>

Siddiqui et al. 1973. Effects of some constant and alternating temperatures on population growth of the pea aphid *Acyrthosiphon pisum* (Homoptera: Aphididae). *The Canadian Entomologist,* 105:1

Campbell and MacKauer. 1975. Thermal constraints for the development of the pea aphid (Homoptera: Aphididae) and some of its parasites. *Cam. Ent.,* 107:4

Özgökçe, M. S., Chi, H., Atlıhan, R., & et al. 2018. Demography and population projection *of Myzus persicae* (Sulz.) (Hemiptera: Aphididae) on five pepper (Capsicum annuum L.) cultivars. *Phytoparasitica*, 46, 153–167. <https://doi.org/10.1007/s12600-018-0651-2>

A. M. Auad, S. O. Alves, C. A. Carvalho, D. M. Silva, T. T. Resende, B. A. 2009. Veríssimo "The Impact of Temperature on Biological Aspects and Life Table of *Rhopalosiphum padi* (Hemiptera: Aphididae) Fed with Signal Grass," *Florida Entomologist*, 92(4): 569-577.

Beetge, L., Krüger, K. 2019. Drought and heat waves associated with climate change affect performance of the potato aphid *Macrosiphum euphorbiae*. *Sci. Rep*., 9: 3645.

<https://doi.org/10.1038/s41598-018-37493-8>

Ölmez, S., Bayhan, E., & Ulusoy, M. R. 2003. Effect of different temperatures on the biological parameters of *Macrosiphum rosae* (L.) (Homoptera: Aphididae). *Journal of Plant Diseases and Protection*, 110(2): 203–208. <http://www.jstor.org/stable/43215525>

TSITSIPIS, J.A. and MITTLER, T.E. 1976. development, growth, reproduction, and survival of apterous virginoparae of *Aphis fabae* at different temperatures. *Entomologia Experimentalis et Applicata*, 19: 1-10. <https://doi.org/10.1111/j.1570-7458.1976.tb02575.x>

Conti, Bruno & Bueno, Vanda & Sampaio, Marcus & Sidney, Livia. 2009. Reproduction and fertility life table of three aphid species (*Macrosiphini*) at different temperatures*. Revista Brasileira de Entomologia*, 54: 654-660. 10.1590/S0085-56262010000400018.

Chang-Gyu Park, Byeong-Ryoel Choi, Jum Rae Cho, Jeong-Hwan Kim, Jeong Joon Ahn. 2017. Thermal effects on the development, fecundity and life table parameters of *Rhopalosiphum padi* (Linnaeus) (Hemiptera: Aphididae) on barley, *Journal of Asia-Pacific Entomology*, 20: 767-775. <https://doi.org/10.1016/j.aspen.2017.05.004>.

Fernandez-Quintanilla, C., Fereres, A., Godfrey, L. and Norris, R.F. 2002. Development and reproduction of *Myzus persicae* and *Aphis fabae* (Hom., Aphididae) on selected weed species surrounding sugar beet fields. *Journal of Applied Entomology*, 126: 198-202.

<https://doi.org/10.1046/j.1439-0418.2002.00627.x>

Feng Hong, Hong-Liang Han, Po Pu, Dong Wei, Jia Wang, Yinghong Liu. 2019. Effects of five host plant species on the life history and population growth parameters of *Myzus persicae* (Hemiptera: Aphididae). *Journal of Insect Science*, 19(5). <https://doi.org/10.1093/jisesa/iez094>

Mala, Mukta & Hemmings, Zac & Andrew, Nigel. (2023). Influence of warming with temperature oscillations on the life history traits of the Aphids *Acyrthosiphon pisum* and *Megoura crassicauda*. 10.21203/rs.3.rs-2817483/v1.

La Rossa, F.R., Vasicek, A. & López, M.C. 2013. Effects of Pepper (Capsicum annuum) Cultivars on the Biology and Life Table Parameters of *Myzus persicae* (Sulz.) (Hemiptera: Aphididae). *Neotrop. Entomol.*, 42: 634–641. <https://doi.org/10.1007/s13744-013-0166-9>

Jiang, Wenbin, Qian Cheng, Changhao Lu, Wenlong Chen, Degang Zhao, and Yingqin He. 2022. "Different host plants distinctly influence the adaptability of *Myzus persicae* (Hemiptera: Aphididae)". *Agriculture*, 12: 2162. <https://doi.org/10.3390/agriculture12122162>

Özgökçe, M.S., Kuşoğlu, D., Konuş, M., Kara, H., Risvanli, M.R., & Çetin, D. 2023. The effects of different Charleston pepper cultivars on the demographic parameters and the antioxidant levels of *Myzus persicae* (Sulzer, 1776) (Hemiptera: Aphididae). *Turkish Journal of Entomology*, 47: 133-147.

del Toro, F.J., Choi, K.S., Rakhshandehroo, F., Aguilar, E., Tenllado, F., & Canto, T. 2019. Ambient conditions of elevated temperature and CO2 levels are detrimental to the probabilities of transmission by insects of a Potato virus Y isolate and to its simulated prevalence in the environment. *Virology*, 530: 1-10. <https://doi.org/10.1016/j.virol.2019.02.001>

Armand, T., Korn, L., Pichon, E., Souquet, M., Barbet, M., Martin, J.-L., Devavry, M., et al. 2021. Efficiency and persistence of Movento® Treatment against *Myzus persicae* and the transmission of aphid-borne viruses. *Plants*, 10(12): 2747. <http://dx.doi.org/10.3390/plants10122747>

Wang et al. 1997. Influence of temperature on development survivorship and reproduction of Buckthorn aphid (Homoptera: Aphididae). *Ann. Entomol. Soc. Am.,* 90: 62-68.

Goundoudaki, S., Tsitsipis, J.A., Margaritopoulos, J.T., Zarpas, K.D. and Divanidis, S. 2003. Performance of the tobacco aphid *Myzus persicae* (Hemiptera: Aphididae) on Oriental and Virginia tobacco varieties. *Agricultural and Forest Entomology*, 5: 285-291. <https://doi.org/10.1046/j.1461-9563.2003.00190.x>

Collins M.C., & Leather, S.R. 2001. Effect of temperature on fecundity and development of the Giant Willow Aphid, *Tuberolachnus salignus* (Sternorrhyncha: Aphididae). *EJE*, 98(2), 177-182. doi: 10.14411/eje.2001.033

Souza MA, Armstrong JS, Hoback WW, Mulder PG, Paudyal S, et al. 2019. temperature dependent development of sugarcane aphids *Melanaphis sacchari*, (Hemiptera: aphididae) on three different host plants with estimates of the lower and upper threshold for fecundity. *Curr Trends Entomol Zool Stds 2*. 1011. DOI:10.29011/CTEZS-1011.001011

Beatriz Maria Diaz and Alberto. 2005. Fereres, Life Table and Population Parameters of *Nasonovia ribisnigri* (Homoptera: Aphididae) at different constant temperatures, *Environmental Entomology*, 34: 527–534, <https://doi.org/10.1603/0046-225X-34.3.527>

Bieri, M., Baumgärtner, J., & Bianchi, G. 1983. Development and fecundity of pea aphid (*Acyrthosiphon pisum* Harris) as affected by constant temperatures and by pea varieties. *Journal of the Swiss Entomological Society*, 56(1-2). <https://doi.org/10.5169/seals-402070>

Nikolakakis, N. N., Margaritopoulos, J. T., & Tsitsipis, J. A. 2003. Performance of *Myzus persicae* (Hemiptera: Aphididae) clones on different host-plants and their host preference. *Bulletin of entomological research*, 93(3), 235–242. <https://doi.org/10.1079/BER2003230>

Kanavaki, O. M., Margaritopoulos, J. T., Katis, N. I., Skouras, P., and Tsitsipis, J. A. 2006.

Transmission of potato virus Y in tobacco plants by *Myzus persicae nicotianae* and *M. persicae*

*s.str*. *Plant Dis.,* 90:777-782.

Erdal Sertkaya and Gulsen Sertkaya. 2005. Aphid transmission of two important potato viruses, PVY and PLRV by *Myzus persicae* (sulz.) and *Aphis gossypii* (glov.) in Hatay province of Turkey. *Pakistan Journal of Biological Sciences*, 8: 1242-1246.
